# Multimodal cues displayed by submissive rats promote prosocial choices by dominants

**DOI:** 10.1101/2022.01.12.475866

**Authors:** Michael Joe Munyua Gachomba, Joan Esteve-Agraz, Kevin Caref, Aroa Sanz Maroto, Helena Bortolozzo-Gleich, Diego Andrés Laplagne, Cristina Márquez

**Affiliations:** Neural Circuits of Social Behaviour Laboratory, Instituto de Neurociencias, Universidad Miguel Hernández - Consejo Superior de Investigaciones Científicas (UMH-CSIC), Sant Joan d’Alacant, Alicante, Spain; Laboratory of Behavioural Neurophysiology, Brain Institute, Federal University of Rio Grande do Norte, Natal, Brazil

**Keywords:** social behaviour, social decision-making, social dominance, prosociality, empathy, behavioural synchrony, ultrasonic vocalisations, body language, rats

## Abstract

Animals often display prosocial behaviours, performing actions that benefit others. Although prosociality is essential for social bonding and cooperation, we still know little about how animals integrate behavioural cues from those in need to make decisions that increase their wellbeing. To address this question, we used a two-choice task where rats can provide rewards to a conspecific in the absence of self-benefit, and interrogated which conditions promote prosociality by manipulating the social context of the interacting animals. While sex or degree of familiarity did not affect prosocial choices in rats, social hierarchy revealed to be a potent modulator, with dominant decision-makers showing faster emergence and higher levels of prosocial choices towards their submissive cage-mates. Leveraging quantitative analysis of multimodal social dynamics prior to choice, we identified that pairs with dominant decision-makers exhibited more proximal interactions. Interestingly, these closer interactions were driven by submissive animals that modulated their position and movement following their dominants and whose 50kHz vocalisation rate correlated with dominants’ prosociality. Moreover, Granger causality revealed stronger bidirectional influences in pairs with dominant focals and submissive recipients, indicating increased behavioural coordination. Finally, multivariate analysis highlighted body language as the main information dominants use on a trial-by-trial basis to learn that their actions have effects on others. Our results provide a refined understanding of the behavioural dynamics that rats use for action-selection upon perception of socially relevant cues and navigate social decision-making.

## INTRODUCTION

Prosocial actions, those that benefit others, support the development of positive social interactions, like cooperation, which sustain individual and group well-being^1–3^. Recent work has demonstrated that prosocial behaviours are not exclusive to humans but conserved across different species^4–24^. Several factors have been proposed to modulate prosocial behaviours, such as familiarity^2, 19, 25^, sex^9, 26, 27^, and social status^28, 29^. Related to the latter, flexibly adapting decision-making based on the social hierarchy of the interacting partner can be cost-effective and, in some cases, a crucial survival strategy. Less effort has been devoted to the identification of the behavioural correlates that lead to such effects, which is important and necessary to determine the proximate mechanisms underlying prosocial choices. Whether enhanced prosociality is due to an improvement in the perception and integration of socially relevant information, or to flexibility in the action-selection process of adequate behavioural strategies upon perception of these social cues, is still far from being understood. Studies at the level of behaviour are needed to identify which are the factors that inform individuals’ social decisions to benefit others.

We evaluated how laboratory rats adapt their decision to help or not to help depending on social context in order to identify the behavioural correlates by which animals incorporate the actions of others into social decision-making. We previously showed that male rats behave prosocially in a two-alternative forced choice task, providing food to a familiar conspecific in the absence of self-benefit, being food-seeking behaviour displayed by recipients necessary for prosociality to emerge ^17^. Here, we used this task to ask about the factors that promote or hinder prosociality by modulating familiarity, sex and social status of the interacting animals. Briefly, our prosocial choice task (PCT) is based on a double T-maze where only the focal animal (decision-maker) controls the access to the food-baited arms of its own and the recipient rat’s maze. In each trial, the focal rat can choose between one side of the maze, providing food only to itself (selfish choice), or the opposite side, providing food to itself and the recipient rat (prosocial choice) (**Figure 1A-C** and **Movie S1**). We hypothesised that social interactions prior to choice might be crucial for increasing the social salience of recipients’ attempts to reach the food and thus might impel decision-makers to learn that their choices have an impact on others. With this aim in mind, we first identified the social conditions where differences in prosociality can be detected and performed a refined analysis of the social interactions observed.

**Figure 1.**
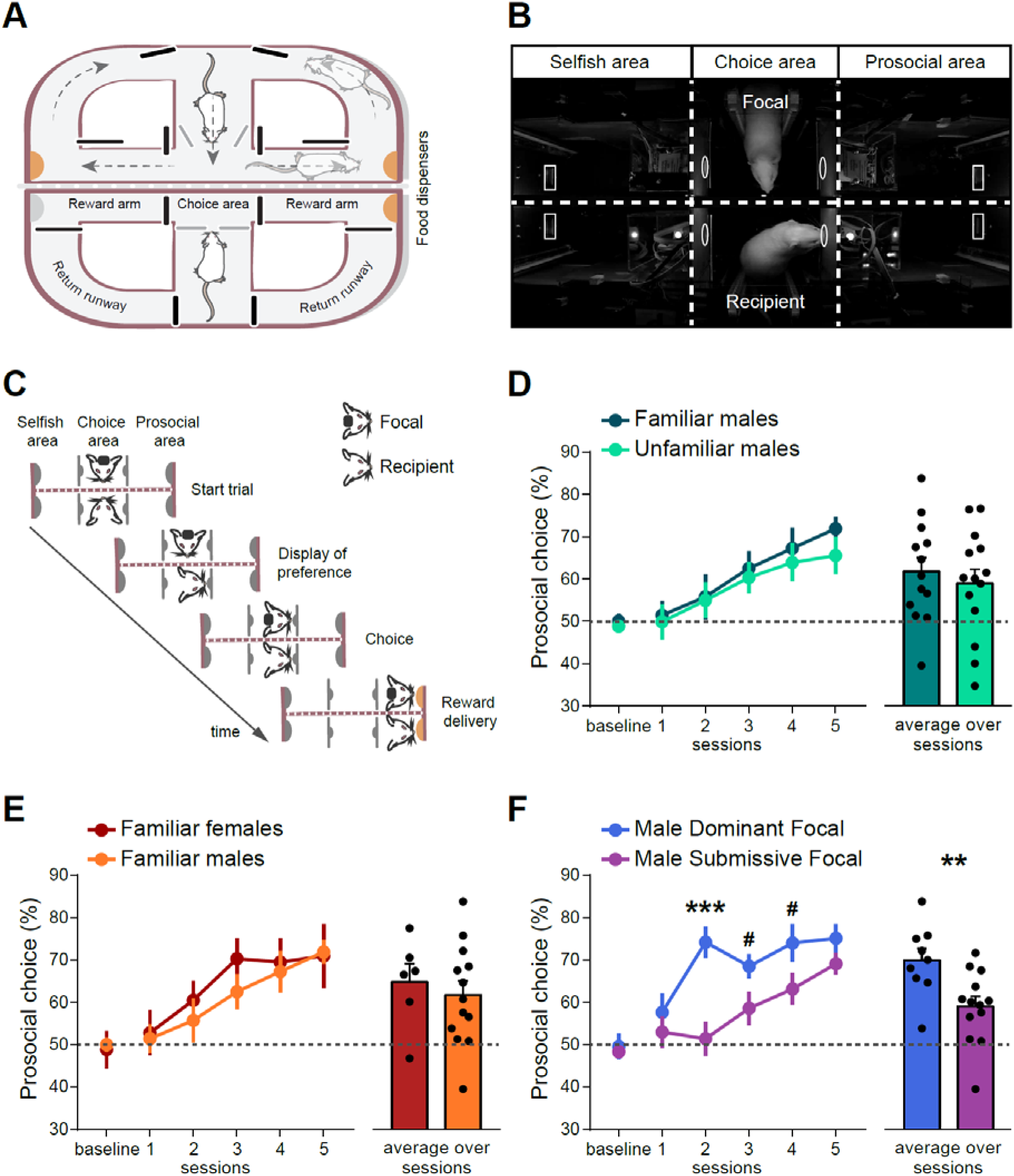
Prosociality emerges faster when decision-makers are dominant, and is not affected by familiarity or sex. (A) Schematic overview of the double T-maze used for the Prosocial Choice Task (PCT). Each T-maze (one per rat) consists of a central arm that gives access to a choice area and two reward areas gated by automated doors (black lines) at the end of which food is delivered (orange semi circles). Access to the choice area is controlled by automated doors placed in the central arm (grey lines). Arrows in the upper maze represent the flow movements of the rats in the maze. (B) Example image from a video recording, showing a top view of the central area of the double T-maze during one session. The horizontal dashed line marks the transparent and perforated wall that separates the two single mazes, which allows rats to see, hear, smell, and partially touch each other. Vertical dashed lines mark the separation between different areas of the maze: the choice area, where social decision-making occurs, and prosocial and selfish areas, where food is delivered depending on the contingencies of the task. Grey ellipses in the choice area mark the position of nose ports, which control the opening of doors located under them. When the decision maker pokes one of its nose ports the door underneath it and the door on the same side for the recipient animal opens, allowing them to reach feeder magazines (grey rectangles in the reward areas). In this example, the focal rat (decision-maker) is in the top of the image, whereas the recipient appears in the bottom, while displaying food-seeking behaviour. (C) Schematic view of a trial: before the PCT focal and recipient rats are trained individually to navigate in the maze and learn to retrieve their own rewards (see Methods). In the PCT, a trial starts when both rats are in the central arm, after opening of the central doors that give access to the choice area. There, the recipient rat will display food-seeking behaviour (repeatedly poking in the side where it was previously trained to find food during individual training), and the focal animal can choose to nose poke on either side of its own maze. Focal animal will always be rewarded; however, recipient’s reward will depend on focal’s choice. A focal’s nose poke on the same side where recipient is displaying food-seeking behaviour (prosocial choice), will lead to both rats receiving one pellet in the reward area, while a nose poke on the opposite side (selfish choice) will lead to only the focal receiving one pellet and the recipient none. Prosocial and selfish sides remain fixed throughout all days, so that the focal animal does not need to read out the behaviour of the recipient on each trial, but can develop a preference over time. After food consumption, rats can pass through the return runway and go back to the central arm to start a new trial. See also **Movie S1**. (D) Familiarity of the interacting animals doesn’t affect prosocial choices in male rats. To understand whether the familiarity of the recipient modulates the proportion of prosocial choices, we compared in the PCT two independent groups: focal animals that performed in the maze with their cage mate (“familiar males”, n=13 pairs) and focal animals that performed with a stranger, non-cage mate, recipient (“unfamiliar males”, n=14 pairs). Unfamiliar animals interacted for the first time in the first session of the PCT, and were maintained over the rest of sessions while not being cage mates. We found that focals of the two groups developed similar proportions of prosocial choices along sessions, indicating that the degree of familiarity of the recipient does not affect prosociality in male rats. (E) Female rats show similar proportions of prosocial choices compared to male rats. To study sex differences in the development of prosocial choices, we tested two independent groups where the focal animal was either male (“familiar males”, n=13 pairs) or female (“familiar females”, n=6 pairs), interacting with a cage mate of the same sex. We found no difference in the proportions of prosocial choices along the five testing sessions, indicating that female and male rats are equally prosocial when interacting with a familiar conspecific of the same sex. (F) Social hierarchy modulates prosocial choices in male rats. Dominant (n=9 pairs) and submissive (n=13 pairs) focals displayed a preference for the prosocial option but dominant focals showed faster emergence and higher proportion of prosocial choices compared to submissive focals. See also **Figure S1**. For D, E, F baseline and five daily test sessions are shown. Baseline corresponds to the percentage of focal’s choices for the side that would later correspond to the prosocial side during testing, averaged across the last two days of individual training. Data represented as mean ± SEM. # p<0.1, **p<0.01, ***p<0.001.

We show that, as observed in non-human primates, male dominant rats are more prosocial with a faster emergence of prosocial biases. Beyond the description of this effect, we unravel the behavioural correlates through which this effect is observed, based on the analysis of social interactions using DeepLabCut for unmarked pose estimation with subsecond resolution, entropy-based algorithms for ultrasonic vocalisation’s agency, granger causality to assess directionality in animal’s interactions and behavioural modelling to identify behaviours predictive of choice on a trial-by-trial basis. Strikingly, dominants’ higher levels of prosociality are a consequence of their submissive partners being better at communicating need and capable of modifying the dominant’s behaviour. This interesting effect emerges in the form of multimodal social dynamics and highlights the importance of embracing the bidirectionality of social interactions in decision making.

## RESULTS

### Prosociality emerges faster when decision-makers are dominant and is not affected by familiarity or sex

We tested pairs of rats in our prosocial choice task (PCT), where a decision-maker rat (focal) can choose in each trial to provide food reward to itself only (selfish choice) or to itself and a recipient rat (prosocial choice) (**Figure 1A-C** and **Movie S1**). After individual training for maze navigation, focal and recipient animals were tested together in PCT, and learned the new reward contingencies where food delivery to the recipient depended on focal’s choices. Each pair performed five daily consecutive sessions of 40 minutes each, over which focals’ choice preference was assessed.

Rats’ prosocial preferences in food-foraging contexts emerged over the testing sessions independently of familiarity or sex. Male rats displayed similar levels of prosociality when interacting with their cage mates or unfamiliar conspecifics (repeated-measures ANOVA with “session” as within-subjects factor and “familiarity” as a between-subjects factor: “session” (F_(4, 100)_ =13.86, p = 5e-9, ƞ^2^=0.164, BF_incl_= 2.961e+6), “familiarity” by “session” (F_(4,100)_ = 0.29, p = 0.882, ƞ^2^=0.003, BF_incl_=0.107) and “familiarity” (F_(1, 25)_ = 0.36, p = 0.555, ƞ^2^=0.008, BF_incl_=0.328)) (**Figure 1D**). Moreover, we did not observe sex differences, with females being equally as prosocial as males (repeated-measures ANOVA: “session” (F_(4, 68)_ = 9.83, p = 2e-6, ƞ^2^=0.181, BF_incl_=71466), “sex and “session” (F_(4, 68)_ = 0.44, p = 0.783, ƞ^2^=0.008, BF_incl_=0.223) and “sex” (F_(1, 17)_ = 0.29, p = 0.596, ƞ^2^= 0.008, BF_incl_=0.391)) (**Figure 1E**). While we did not find evidence for an effect of familiarity or sex in prosocial tendencies, it could be that the proportion of prosocial individuals would differently emerge over the testing sessions in each group. For this, we computed a Prosocial Choice Index (PCI) that reflected the strength of the prosocial (or selfish) bias compared to chance. Using a permutation test we categorized the animals as either prosocial, unbiased or selfish over the days. The emergence of prosociality was comparable across groups (**Figure S1A-F and Supplementary Table 1**).

To understand how social dominance may modulate prosocial choice, we first identified the social status within pairs of cage mate rats. For this, we used the modified Food Competition Test^30^, a novel trial-based dominance assay, where established social hierarchies can be identified in the home cage of non-food deprived pairs of male rats. It has the added advantage of not inducing aggressive interactions during testing which could influence later prosocial tendencies. After identification of social status of the animals (**Figure S1J**), we tested for prosocial tendencies two parallel independent groups, where the decision-maker rat was either the dominant (and thus its recipient was submissive) or the submissive (and the recipient was the dominant). Thus, in both groups a dominant animal would interact with a submissive, but their roles in the decision process would differ. We found that both groups acquired a preference for the prosocial option over the days but that social hierarchy drastically modulated the emergence of this choice (**Figure 1F**). Specifically, dominant animals acquired faster prosocial tendencies and reached higher prosociality levels than submissive decision-makers (repeated-measures ANOVA: “session” (F_(4, 80)_=8.42, p=1e-5, ƞ^2^=0.15, BF_incl_= 3445), “hierarchy” by “session”(F_(4,80)_=2.67, p=0.038, ƞ^2^=0.048, BF_incl_= 5.8) and “hierarchy”(F_(1, 20)_=8.75, p=0.008, ƞ^2^=0.136, BF_incl_=11.2)). Dominant and submissive decision-makers displayed similar choices on the first session of the prosocial choice task, where animals are exposed to the social task for the first time after individual training and had not yet learned that their actions have consequences on the reward contingencies of the recipient (*t* test of the proportion of prosocial choices of dominant focals against submissive focals: t_(20)_ = 0.81, p = 0.428, BF_10_=0.491 for day 1). However, marked differences appeared from the second day of testing, where dominant animals displayed strong prosocial preferences while submissive focals were still at chance levels (independent sample *t* test t_(20)_ = 4.03, p = 6.5e-4, BF_10_=42, for day 2; paired sample *t* test of proportion of prosocial choices in session 2 against baseline for dominant focals t_(8)_ = 5.72, p = 4.4e-4, BF_10_=80; for submissive focals t_(12)_ = 0.681, p = 0.509, BF_10_=0.34). Interestingly, prosociality on this day positively correlated with the strength of the social hierarchy (**Figure S1L**) suggesting a parametric relationship between dominance and prosociality (Pearson correlation between prosocial choice in day 2 and Dominance Index r=0.71, p < 0.001). The differences between dominant and submissive focals were maintained over the sessions but progressively faded once submissive focals started to show prosocial biases from day 3 onwards (t test t_(20)_ = 1.87, p=0.077, BF_10_= 1.28, for day 3; t_(20)_ = 1.88, p = 0.074, BF_10_=1.3, for day 4; t_(20)_ =1.42, p = 0.171, BF_10_=0.79, for day5). We then assessed whether the proportion of prosocial, unbiased and selfish animals would be different depending on their hierarchical status and observed a higher proportion of prosocial animals in pairs with dominant focals in the second day of testing (**Figure S1G-I**).

While previous non-human primate studies also showed evidence of prosociality occurring down the hierarchy, the factors leading to such directionality are not known. Leveraging the controlled environment that experiments in laboratory rats provide, we endeavoured to identify the behavioural correlates at the base of this enhanced prosociality in dominant animals. To this end, we performed a fine-grained analysis of rats’ behaviour during the choice period (time from trial start to focal’s choice), focusing our analyses on the first two days of the task, when prosocial bias emerges, in order to identify the behavioural dynamics that promote integration of actions from others into decision making processes.

### Social dominance does not affect recipients’ food-seeking behaviour nor focals’ latency to decide

Previously, we demonstrated that the recipient’s display of food-seeking behaviour –poking in the nose port that gives access to the food-baited arm– is necessary for the emergence of prosocial choices by focal rats^17^. Thus, one possibility was that submissive recipients were better at displaying food seeking behaviour, facilitating the learning of the contingencies of the task by dominant decision-makers. However, we did not find hierarchy differences in the number of nosepokes performed, nor on the vigour with which they were displayed (**Figure S2A**). Dominant humans are faster in making (non-social) decisions in stressful situations, without compromising their accuracy^31^. However, we did not observe differences in the latency to choose in rats performing our task (**Figure S2B**). It could still be possible that dominant focals developed faster prosocial preferences in the first days of testing because of increased task performance, thus accelerating the learning rate of the new contingencies in the social task. However, this was not the case either (**Figure S2C**). We then hypothesised that the social interactions displayed prior to the choice might be at the core of the faster learning of contingencies for dominant focals, and that those pairs with dominant decision-makers would display richer social interactions.

### Social dominance modulates the quality but not the quantity of social interactions prior to choice

We analysed trial-by-trial social interactions in off-line video recordings using Bonsai^32^ and DeepLabCut^33^ which enabled us to precisely extract the position of unmarked body-parts of the interacting animals with high spatial and temporal resolution. The time animals spent directly investigating each other was equivalent regardless of the pronounced differences in prosociality (**Figure 2A,** independent sample t test for “mutual direct investigation”: t_(20)_=0.411, p=0.685, BF_10_=0.413). Although direct contact is the standard measure of social interaction, we hypothesised that significant social interactions might still happen at a distance, and not only through direct sniffing of the partner. Thus, we quantified the time that animals spent simultaneously in the choice area, regardless of the distance between them. Again, no differences were observed on the duration of these distant social interactions according to social status (**Figure 2B,** t_(20)_=0.047, p=0.96, BF_10_=0.39).

**Figure 2.**
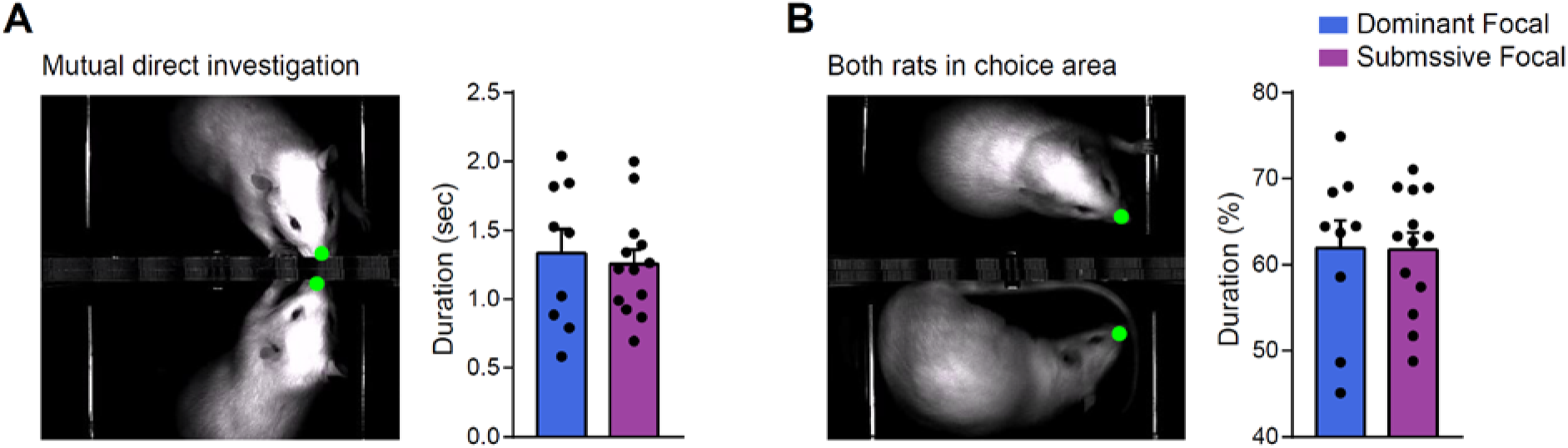
Social dominance does not affect the quantity of social interactions prior to choice. (A) Social dominance does not affect the duration of social investigation prior to the choice nor (B) the percentage of choice time per trial that both rats are present in the choice area, as an index of social interactions in the distance. Data is shown as mean ± SEM, individual dots show the averaged trial value for each pair across the first two sessions.

Even though the duration of mutual direct investigation and interaction time during choice was similar in the two groups, it was still possible that dominance status could account for differences in the quality of the social dynamics when animals were at a social distance. To this end, we quantified on each trial the median value of the distance between the focal and the recipient rat while in the choice area, referred to as nose-to-nose distance, as a measure of social proximity. Indeed, pairs with a dominant focal maintained a closer distance prior to choice (independent sample *t* test: t_(20)_ = -2.53, p = 0.020, BF_10_=3.15, **Figure 3A**). Interestingly, this effect was already present in the first testing session, where no differences in prosocial choice were yet observed (**Figure S3A-C**). Pairs with dominant decision-makers displayed closer interactions in a higher proportion of trials (**Figure 3B**), and these differences emerged early during the interaction time, where pairs with a dominant decision-maker would be closer to each other than those with a submissive focal, when the focal was going to choose the selfish option (**Figure 3C** left panel; two-way ANOVA, “hierarchy” by “choice” F_(1, 1986)_ = 4.77, p = 0.029; “choice” F_(1,1986)_ = 294.5, p = 1e-61 and “hierarchy” F_(1,1986)_ = 13.89, p = 0.0002; further decomposition of the interaction followed by SNK post hoc test revealed a significant difference across dominance categories in selfish but not prosocial trials F_(3, 1986)_ = 120.3, p = 1e-71, **Figure 3C** right panel).

**Figure 3.**
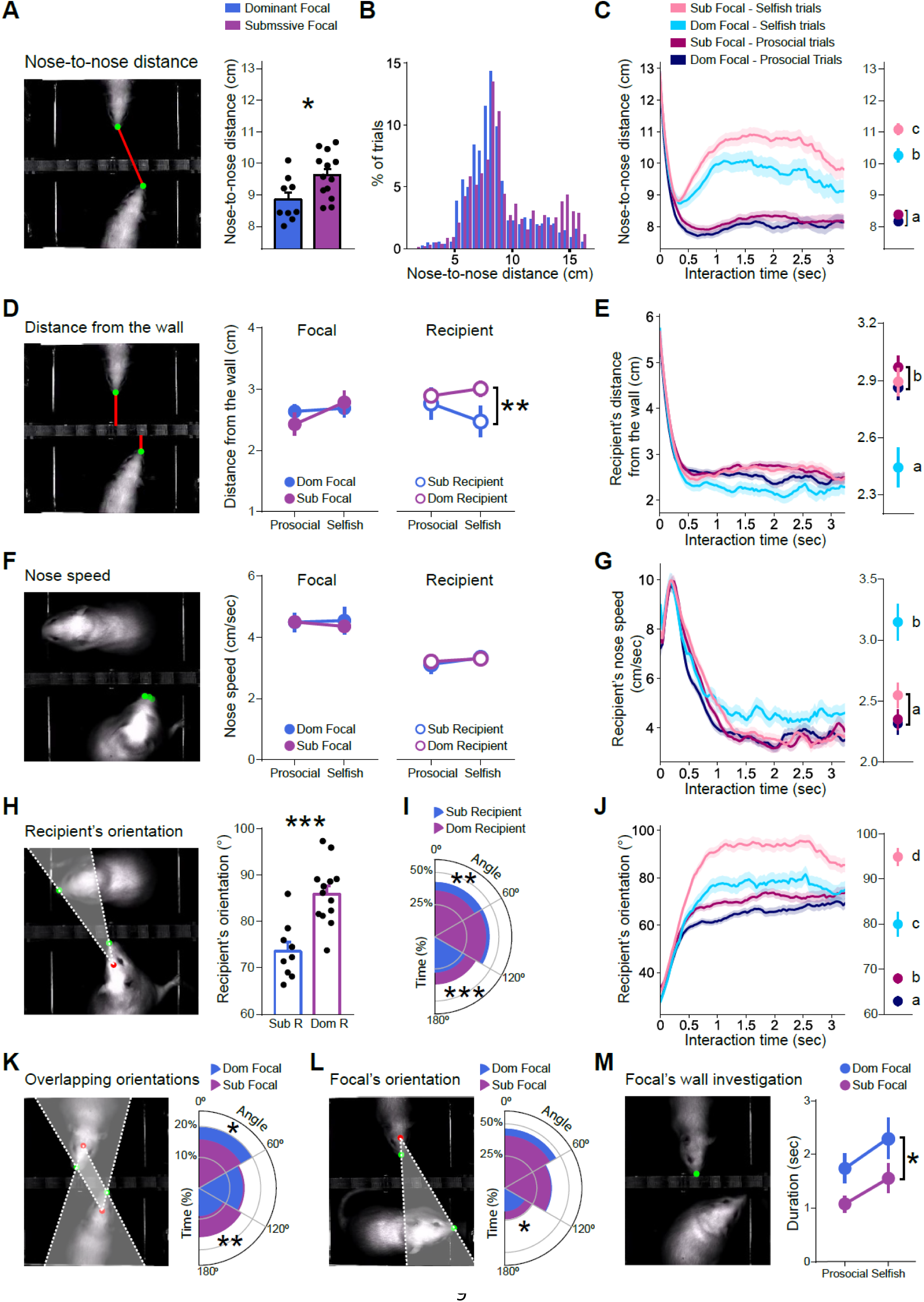
Social dominance modulates the dynamics of social interactions prior to choice. (**A-C**) **Pairs with dominant rat as focal display more proximal interactions prior to choice.** (**A**) The distance between focal and recipient noses, as a proxy for social interest of the pair, was measured during the interaction time, defined as the time that the two rats were simultaneously present in the choice area. The median nose-to-nose distance per trial was lower in pairs with dominant focals. Moreover, (**B**) the proportion of trials with closer interactions was higher when dominants were decision-makers, being (**C**) these more proximal interactions already evident in the first seconds of interaction and only observed in selfish trials (see **left panel** showing temporal dynamics; **right panel** showing the average, SEM and statistics of this time window). (**D-G**) **Submissive recipients follow their dominant decision-makers.** (**D**) To identify if one of the interacting animals was driving these more proximal social interactions, we measured the distance between the nose of each rat and the dividing wall that separated the animals in the choice area. The median distance from the wall per trial was similar for focal rats across dominance categories and trial type, while a tendency was found for submissive recipients to stay closer to the wall in selfish trials, suggesting an increased social interest towards their dominant focals when they were going to choose not to reward them. (**E**) This tendency was present during the early phase of interaction. (**F**) Movement dynamics indicated a similar pattern, where median nose speed in the whole choice period did not revealed differences but (**G**) submissive recipients showed higher nose speeds during the first seconds of interaction in selfish trials, suggesting again that they were following their dominant when it was going to poke in the selfish side. (**H-L**) **Pairs with dominant focal and submissive recipient display more coordinated gazing.** Orientation of each animal towards the partner was calculated as the angle between the vector from the centre of its head (red dot) to its own nose (green dot), and the vector from the red dot and to the partners’ nose. Lower values indicate more directed gazing. (**H**) Submissive recipients were more oriented towards their focal when considering median head orientation towards the nose of the partner per trial, regardless of trial type, and (**I**) spent a higher proportion of time directly oriented in angles smaller than 60°. (**J**) These differences in head orientation were evident in the first seconds of interaction, being submissive recipients more oriented towards their dominant focal both in prosocial and selfish trials. (**K**) Pairs with dominant focal spent a higher proportion of time orienting to each other, while pairs with submissive focal spent a higher proportion of time orienting away from each other. (**L**) The same tendency was observed in the head orientation of the focals, although only significant in the case of submissive focals, which spent a higher proportion of time orienting away from their recipient, compared to dominant focals. (**M**) Dominant focals spent more time investigating the wall when their submissive recipient was in the choice area, compared to submissive focals, both in prosocial and selfish trials, indicating a higher attention towards their recipients’ behaviour and suggesting increasing sniffing through the wall. MEAN ± SEM is shown. *p<0.05, **p<0.01, ***p<0.001. In C, E, G, J right panels: letters denote statistically significant differences between conditions with significant level set to 0.05. Sub= subordinate; Dom= dominant; R= recipient. See also **Figure S3** with analysis by days.

Social interactions are by definition bidirectional and highly dynamic^34^ and although classical studies on decision making have focused on the analysis of the decision-maker, it could well be possible that focals were influenced by the behaviour of the recipient animal. To ascertain which animal (focal or recipient) was responsible for these more proximal interactions, we quantified the median distance between each rat’s nose and the central wall that divided the two mazes, as a proxy for social interest (**Figure 3D** and **S3D-F**). No significant differences were found between dominant and submissive decision-makers (**Figure 3D** middle panel, repeated-measure ANOVA, “choice” (F_(1,20)_ = 7.78, p = 0.011, BF_incl_=6.49), “choice” by “hierarchy” (F_(1,20)_ = 4.32, p = 0.051, BF_incl_=1.76) and “hierarchy” (F_(1,20)_ = 0.05, p = 0.820, BF_incl_=0.58)). However, submissive recipients were closer to the wall on selfish trials compared to dominant recipients (**Figure 3D** right panel, “choice” (F_(1,20)_ = 2.027, p = 0.170, BF_incl_=0.37), “choice” by “hierarchy” (F_(1,20)_ = 10.86, p = 0.004, BF_incl_=8.44) and “hierarchy” (F_(1,20)_ = 1.76, p = 0.200, BF_incl_=0.87); independent sample *t* test for recipient rats in selfish trials: t_(20)_ = -1.859, p = 0.088). Further analysis showed that this tendency for submissive recipients to stay closer to the wall was occurring during the early phase of interaction when decision-makers were going to choose to be selfish (**Figure 3E** left panel; two-way ANOVA, “hierarchy” by “choice” (F_(1, 1986)_ = 5.38, p = 0.020), “choice” (F_(1, 1986)_ = 11.17, p = 0.001) and “hierarchy” (F_(1, 1986)_ = 14.26, p = 0.0001); further decomposition of the interaction : (F_(3, 1986)_ = 7.388, p=6e-5) followed by SNK post hoc test revealed that the distance from the wall of submissive recipients in selfish trials was different from the other three categories, **Figure 3E** right panel).

The above results indicate that dominance status affects the recipient’s behaviour: submissive recipients stay closer to the wall during selfish trials, thus decreasing the distance from the focal rat. Since, in principle, the act of nosepoking would lead rats to show similar nose movements and trajectories, the increased proximity of submissive recipients to the wall may suggest that these animals move and/or orient themselves towards the focal when it is going to choose the selfish poke. Indeed, analysis of animals’ movement (**Figure 3F** and **S3G-I**) showed that recipient rats continued moving the snout when the trial was going to be selfish. Similar values of nose speed were found across dominance categories for both focal and recipient rats (**Figure 3F**; no significant effects were found for focal nor recipient rats, repeated-measure ANOVA: “choice” (focals: F_(1,20)_ = 0.07, p = 0.798, BF_incl_=0.31; recipients: F_(1,20)_ = 2.526, p = 0.128, BF_incl_=0.71), “choice” by “hierarchy” (focals: F_(1,20)_ = 0.34, p = 0.568, BF_incl_=0.44; recipients: F_(1,20)_ = 0.354, p = 0.559, BF_incl_=0.43) and “hierarchy” (focals: F_(1,20)_ = 0.06, p = 0.814, BF_incl_=0.51; recipients: F_(1,20)_ = 0.017, p = 0.898, BF_incl_=0.55). Nevertheless, the dynamics of nose speed in the early phase of interaction (**Figure 3G** left panel) showed that nose speed of submissive recipients was higher on selfish trials, especially after the first second of interaction (**Figure 3G** right panel; two-way ANOVA, “hierarchy” by “choice” (F_(1,1925)_ = 8.76, p =0.003), “choice” (F_(1,1925)_ = 22.93, p = 2e-6) and “hierarchy” (F_(1, 1925)_ = 6.96, p = 0.008); one-way ANOVA dissecting the interaction (F_(3, 1925)_ = 9.609, p = 0.000003) followed by SNK post hoc test, nose speed of submissive recipients in selfish trials was higher compared to the other three categories).

We further asked whether dominance status affected the degree to which recipient rats were orienting towards their focal, as indication of increased social attention. To this end, we measured the orientation angle of the recipient’s head towards the focal nose (**Figure 3H** left panel), and found that submissive recipients were more oriented towards their focal compared to dominant recipients, with lower values indicative of a more directed orientation (**Figure 3H** right panel; independent sample *t* test t_(20)_ = -4.52, p = 0.0002, BF_10_=106.48). Further analyses revealed that over the interaction time prior to choice, submissive recipients spent a higher proportion of time orienting towards their focal, while dominant recipients spent a higher proportion of time orienting away from their focal (**Figure 3I**; independent sample *t* test for the proportion of time with orientation <60°: t_(20)_ = 3.80, p = 0.001, BF_10_=27.42, time with orientation between 60° and 120°: t_(20)_ = 1.18, p = 0.253, BF_10_= 0.63, time with orientation >120°: t_(20)_ = - 4.32, p = 0.0003, BF_10_=72.84). The same effect was observed in the dynamics of orientation during the early phase of interaction (**Figure 3J** left panel), with submissive recipients more oriented to their dominant decision-maker both in prosocial and selfish trials (**Figure 3J**, right panel; two-way ANOVA, “hierarchy” by “choice” (F_(1,1985)_ = 9.20, p = 0.002); “choice” (F_(1,1985)_ = 183.43, p = 4e-40) and “hierarchy” (F_(1,1985)_ = 37.56, p = 1e-9); one-way ANOVA dissecting this interaction (F_(3,1985)_ = 91.87, p = 1.136e-55), followed by SNK post hoc test revealed a significant difference between all conditions). These results suggest that submissive recipients are more attentive to the behaviour of the focal rat before the decision and change their orientation and position to maintain closer interactions with their dominant partner. Interestingly, this increased gazing from submissive recipients towards their dominant decision-maker was already observed in the first day of testing, while prosocial biases were not yet present (**Figure S3J-M**).

Next, we assessed whether this behaviour of the recipient would lead to a more coordinated reciprocal interaction. We found that indeed, over the interaction period, pairs with dominant focals spent a higher proportion of time orienting to each other, while pairs with submissive focals spent a higher proportion of time orienting away from each other (**Figure 3K**, independent sample *t* test for the proportion of time with both rats orientations <60°: t(20) = 2.36, p = 0.029, BF_10_=2.43, both rats orientations between 60° and 120°: t(20) = -0.33, p = 0.742, BF_10_=0.4, time with both rats orientations >120°: t(20) = -3.37, p = 0.003, BF_10_=12.55). In addition, submissive focals spent a higher proportion of time orienting away from their recipient, compared to dominant focals (**Figure 3L**, independent sample *t* test focal orientation <60°: t_(20)_ = 1.50, p = 0.148, BF_10_=0.85; from 60°-120°: t_(20)_ = 0.34, p = 0.739, BF_10_=0.40; orientation >120°: t_(20)_ = -2,42, p = 0.025, BF_10_=2.66). Importantly, although orientation of focal animals was not as strongly modulated by hierarchy as observed for recipient’s or mutual orientations, dominant decision-makers spent more time directly sniffing through the wall during the interaction period suggesting enhanced social interest (**Figure 3M**; repeated-measure ANOVA, “choice” (F_(1,20)_ = 5.70, p = 0.027, BF_incl_=3.29), “choice” by “hierarchy” (F_(1,20)_ = 0.04, p = 0.844, BF_incl_=0.35) and “hierarchy” (F_(1,20)_ = 4.95, p = 0.038, BF_incl_=2.16)). This effect was mainly driven by the behaviour displayed in the second day of testing, when prosociality emerged (**Figure S3N**), and was not observed in recipient animals (**Figure S3O**).

### Granger causality analysis of focal and recipient movements in the choice area reveals increased bidirectional influence in pairs with dominant focal

Overall, the results so far suggest that submissive recipients are more attentive to their dominants: they display more direct gazing prior to choice and increase proximity to their focals, specifically when decision-makers are going to be selfish (i.e., following them around the choice area). Dominant decision-makers might respond to these cues by showing increased social attention to their recipients which is reflected in increased sniffing time directed to the animal that needs help. In order to establish directionalities in the interactions between focals and recipients within trials we implemented Granger Causality from Partial Directed Coherence, that evaluates whether the past of one time series contains exclusive information that helps predict the present value of another one. We computed the position of each rat’s nose along the x axis (parallel to the dividing wall and ranging from the selfish port to the prosocial port) as a proxy for body movement between the two choice options (**Figure 4A**), assessed whether the position of a rat would cause the other to follow (or move away), and whether this was dependent on hierarchy. Indeed, dominant focals strongly granger-caused (g-caused) the position of their submissive recipient (**Figure 4B**, 0.018 bits, p = 0.002 against trial-shuffled surrogates, see methods) and vice versa (**Figure 4C**, 0.006 bits, p = 0.001) indicating that both animals g-caused changes in the position of the other (**Figure 4D**). Considering that the positions of the animals are positively correlated (Pearson r=0.277, p<0.00001, n=20180), these results suggest that the movement of an animal causes movement of the other in the same direction, compatible with following behaviour. The position of the rats was also positively correlated in dyads with submissive decision-makers (Pearson r=0.146, p<0.00001, n=26628), however, we found causality from focals to recipients (**Figure 4E**, 0.003 bits, p = 0.007) but not from recipients to focals (**Figure 4F**, 0.001 bits, p = 0.060), that is, unidirectional transfer of information (**Figure 4G**).

**Figure 4.**
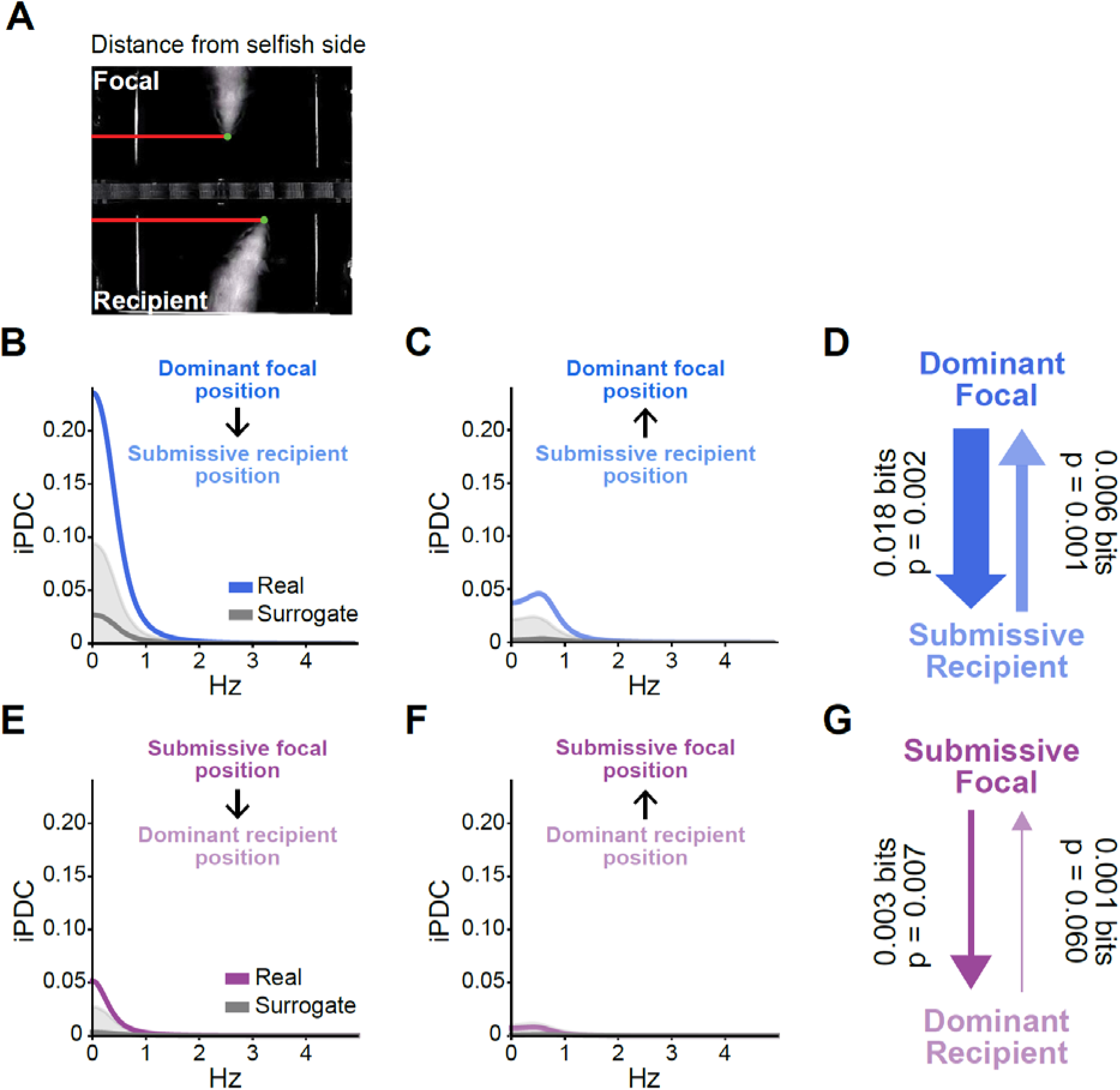
Granger Causality analyses of animals’ position reveal stronger bidirectional influences in dyads with dominant focals. (**A**) Rats’ position was measured as horizontal distance from the selfish side of the choice area. (**B**) Information Partial Directed Coherence (iPDC) from dominant focal to submissive recipient and (**C**) from submissive recipient to dominant focal. (**E**) iPDC from submissive focal to dominant recipient and (**F**) from dominant recipient to submissive focal. iPDC spectra from the real data are shown together with median and 95% confidence intervals from surrogate spectra distributions. (**D**) Information flow (Iflow) representing the causality from dominant focal to submissive recipient and vice versa in units of information transfer. (**G**) Iflow from submissive focal to dominant recipient and viceversa. Arrow widths are proportional to the Iflow values in each direction. P values account for significant differences between the real and surrogate iPDC.

Interestingly, although both decision-makers g-caused changes in the position of their recipients, this influence was stronger when focals were the dominant of the pair (contrast Focal Dominant to Recipient Submissive larger than Focal Submissive to Recipient dominant, p=0.006). Importantly, the influence that the movement of the recipients caused in their decision-makers was stronger in the case of submissive recipients (contrast Recipient Submissive to Dominant Focal larger than Dominant Recipient to Submissive Focal, p=0.032). Altogether, these results indicate that decision-maker and recipient become interdependent by influencing each other’s movements, with dominance affecting the strength of such increased coordination.

### Social dominance modulates recipient’s call rate prior to choice which correlates with the emergence of prosociality

In addition to body position, movement and orientation, rats exchange social information through acoustic signals^35, 36^. Adult rats emit vocalisations in ultrasonic frequencies of two distinct families: the 22-kHz or “alarm calls” and the 50-kHz calls^36^. The latter have been linked to different features of rat behaviour, including mating^37, 38^, play^39^, social contacts^40^, reward anticipation^41^, sniffing and locomotor activity^42–44^. However, the role of USVs as communicative signals mediating animal prosocial decision-making has been largely unexplored. To address this, we recorded USVs during the first two days of the prosocial choice task, performed automated assignment of USVs agency based on the entropy of the signal (**Figure 5A, Figure S4A-B**) and combined this information with tracking and behavioural data in the maze (**Figure 5B-C and Movie S2**) in order to examine how this multimodal information may relate to dominance status and the emergence of prosocial choices.

**Figure 5.**
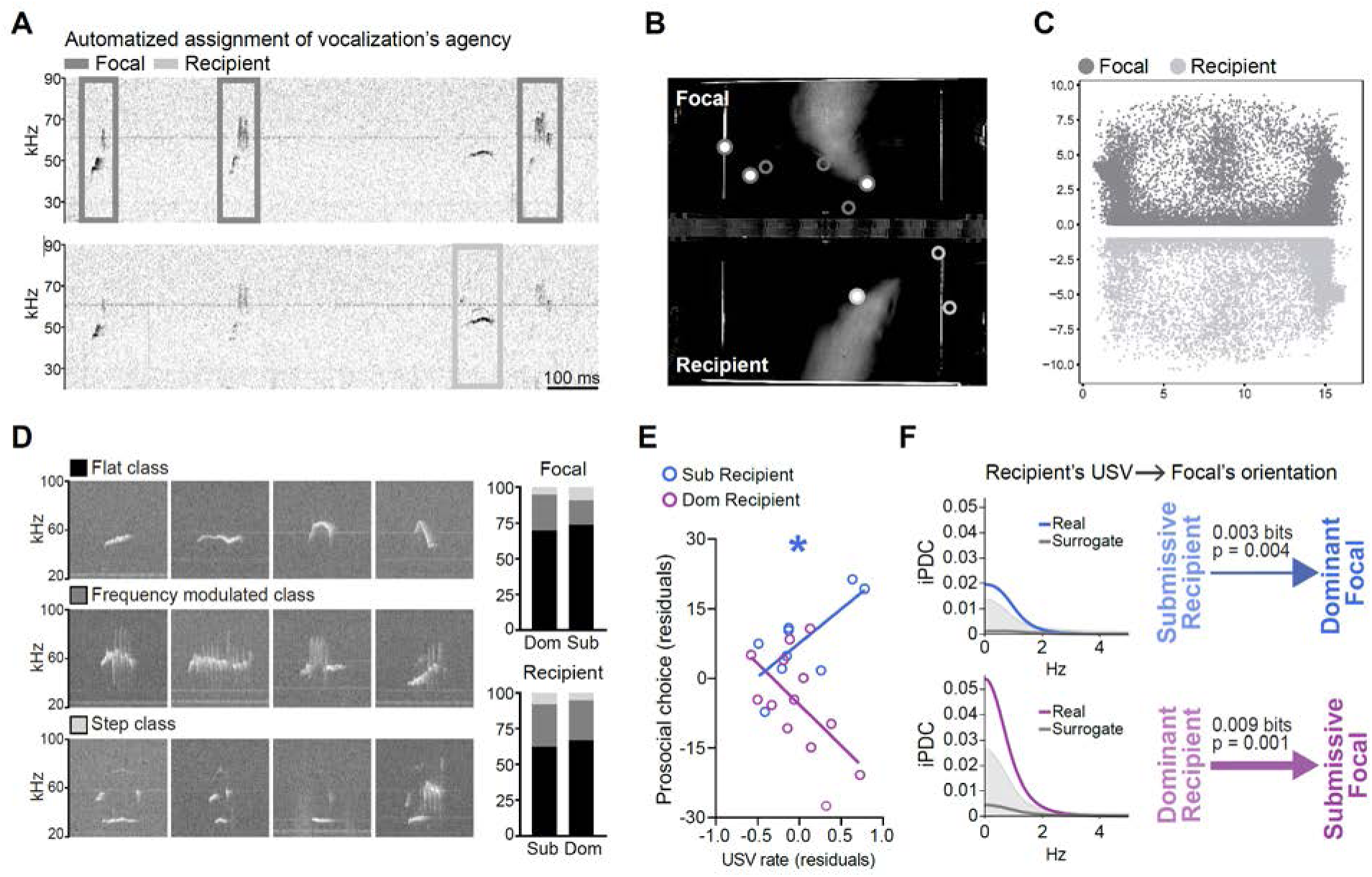
Social dominance modulates recipient’s call rate prior to choice which correlates with the emergence of prosociality. (**A**) Example USVs recording from a prosocial choice task session, showing sonograms for the two microphones, each one placed above the choice area of each maze. In this case, the top sonogram corresponds to the microphone placed above the focal rat and the bottom one to that on top of the recipient. Notice that USVs are detected from both microphones but each USV is automatically assigned to either the focal (dark grey rectangles) or the recipient rat (light grey rectangles) according to the entropy levels (see methods). (**B**) Example image showing localization of agent-assigned USVs emitted during a trial by the focal and recipient rats while in the choice area. Circles indicate the position of the rats’ nose at the time a USV was emitted. Filled circles correspond to the USVs shown in (A). (**C**) Nose location of focal (dark grey) and recipient (light grey) for all USVs detected, relative to the choice area, during the first two days of the prosocial choice task. USVs were emitted in the whole choice area, however they were more frequent around nose-pokes and in proximity to the wall that separated both mazes. See also **Figure S4**. (**D**) Four examples of spectrogram images are shown for each USV class: flat, frequency modulated and step. Flat calls were the most frequent class observed, followed by frequency modulated, while step calls were rare. The proportions of calls (right panel) were similar in focals (top) and recipients (bottom) regardless of the hierarchy status (see **Figure S5**). (**E**) Partial correlation between recipients’ USVs rate and focals’ prosocial choice preference in the first two days of the PCT, when recipients’ speed was regressed out, indicating that the more the submissive recipient vocalises, the more prosocial their dominant partner would be. This correlation was only marginally significant when recipients were the dominant in the pair. (**F**) Granger causality from recipients’ USV to focals’ orientation, showing Information Partial Directed Coherence (iPDC) from submissive recipient to dominant focal (top) and from dominant recipient to submissive focal (bottom). Independently of the hierarchy status of the animals, recipients’ USVs would granger-cause an orientation response from the focal rat. Real iPDC values, surrogate median and 95% confidence intervals of the surrogates’ distribution are shown. Arrow widths are proportional to the Iflow and p-values account for significant differences between the real and surrogate iPDC. *p<0.05.

All USVs recorded during the task were of the 50-kHz family –i.e., no alarm calls were observed– suggesting a positive emotional state of the interacting rats. Many vocalisations were emitted when the nose of the rats was close to the wall separating the two individual mazes and around the nose-ports (**Figure 5C**). Nevertheless, normalising the call rate by nose location revealed that rats vocalised with similar rates throughout the choice area, with no clear spatial preference (**Figure S4C**). Consistent with previous findings^44^, rats in our task (both focal and recipient) partially synchronized the emission of calls with their own body movement, as evidenced by temporally precise correlations between nose speed and vocal production (**Figure S4D-E**). Interestingly, call rate was specifically modulated according to the role each animal had in the task, where focal animals vocalised at a higher rate than their recipients (**Figure S4D**).

To explore whether there were qualitative differences in the calls emitted by the animals, we classified their vocalisations into three different classes corresponding to different vocal programs (flat, frequency modulated and step class). For this, we used VocalMat^45^, a novel platform using convolutional neural networks for sonogram-based classification of rodent USVs. We did not find differences on the qualitative nature of USVs in focal/recipient or dominant/submissive animals (repeated-measures ANOVA with “USV class” as within-subjects factor and “hierarchy” as between-subjects factor. For focal rats: “USV class” (F(1.36, 27.26)= 100.57, p = 7e-12), “USV class” by “hierarchy” (F(1.36, 27.26)=0.05, p = 0.322) and “hierarchy” (F(1,20)=0.01, p = 0.912); for recipients: “USV class” (F(1.07, 21.35)= 80.38, p = 7e-9), “USV class” by “hierarchy” (F(1.07, 21.35)=0.36, p = 0.567) and “hierarchy” (F(1,20)=0.04, p = 0.847) (**Figure 5D**), neither in the evolution of this proportion across days and trial type (**Figure S5A-C**).

Then, we asked how focals’ prosocial choices were related to the vocalisation rates of the interacting animals. We included nose speed of the emitting rat as cofactor to isolate specific modulations of USVs rates from possible variations in movement. Prosocial choices were positively correlated with recipient’s call rate but only when the recipient was the submissive of the pair (partial correlation between USV rate and prosocial choices, controlling for recipient speed: for submissive recipients r = 0.73, p = 0.037; for dominant recipients: r = -0.56, p = 0.055) (**Figure 5D and Figure S5D**). Since submissive recipients were also found to modulate their position and movement towards the focal, these results suggest that they may increase call rate to interact further with the focal, consistent with the proposed role of 50-kHz calls in promoting social contact^46^.

Interestingly, we found that social hierarchy modulated the direction of correlations between USV rate and prosociality where, the more submissive rats would call the more prosociality, and conversely the more dominants would call the less prosociality would be observed, especially on the second day of testing (**Figure S5D**). Although the sign of the correlations was independent of the role (focal/recipient) in the task, these correlations were mainly significant when considering the USV rate of recipients.

In light of this opposite correlation, we asked whether the effect that recipients’ calls have on their decision-makers’ behaviour was different depending on their hierarchical status. We used Partial Directed Coherence to test whether the emission of USVs by submissive and dominant recipients would affect gazing behaviour of their focal differently (**Figure 5F**). We found that emission of calls from the recipient rat promoted more direct gazing from the focal, but this was independent of social hierarchy (I_flow_ from submissive recipients’ USV to dominant focals’ orientation: 0.003 bits, p = 0.004; Pearson r=-0.0732, p=7e-32 (n=25641); I_flow_ from dominant recipients’ USV to submissive focals’ orientation: 0.009 bits, p = 0.001; Pearson r=-0.0528, p = 5e-21 (n=31706)).

### Identification of multimodal cues displayed by both animals as predictors of prosocial choices on a trial-by-trial basis

So far, we described that dominant animals are more prosocial, learning the contingencies of the prosocial choice task faster, and submissives, by following their dominants, have a stronger impact when communicating need. This is related to a more synchronised social interaction of both animals that builds upon multimodal cues displayed by submissive recipients, especially when decision-makers are going to behave selfishly. These different social dynamics are correlated with prosocial choice, however it is still uncertain which cues animals utilise that predict prosocial choices on a trial-by-trial basis. To examine the contribution of a multitude of parameters to the focal’s choice, we employed a multi-step Generalised Linear Model (GLM). Given that nose-to-nose distance and the gazing angle of each animal of the pair were the regressors that explained most of the deviance (see methods, **Figure 6A-B** and **Figure S6**), we asked how these parameters interact with social hierarchy and trial progression in the prediction of prosociality. Trial progression, often included in models of decision-making as a proxy for learning, was considered critical for our analyses as animals starting the social task learn about the new contingencies with respect to the individual training. Significant interactions between behavioural variables, learning and social hierarchy were observed (**Supplementary Table 2**). As our main objective was to disentangle the contribution of social hierarchy on the predictors of choice, we performed reduced GLM for data from each dominance category (**Figure 6C-D**). In both social hierarchy groups, trial progression positively influenced prosocial choice, i.e., both groups increased prosocial choice over time. However, only when dominant animals were the focals were there additional behavioural changes as sessions proceed. Specifically, when dominant animals were the decision-makers, orientation angles decreased (i.e., there was more direct gazing) as trials proceeded, indicating the occurrence of some form of learning on behalf of both animals that ultimately led to a higher proportion of prosocial choices (**Supplementary Table 3**).

**Figure 6.**
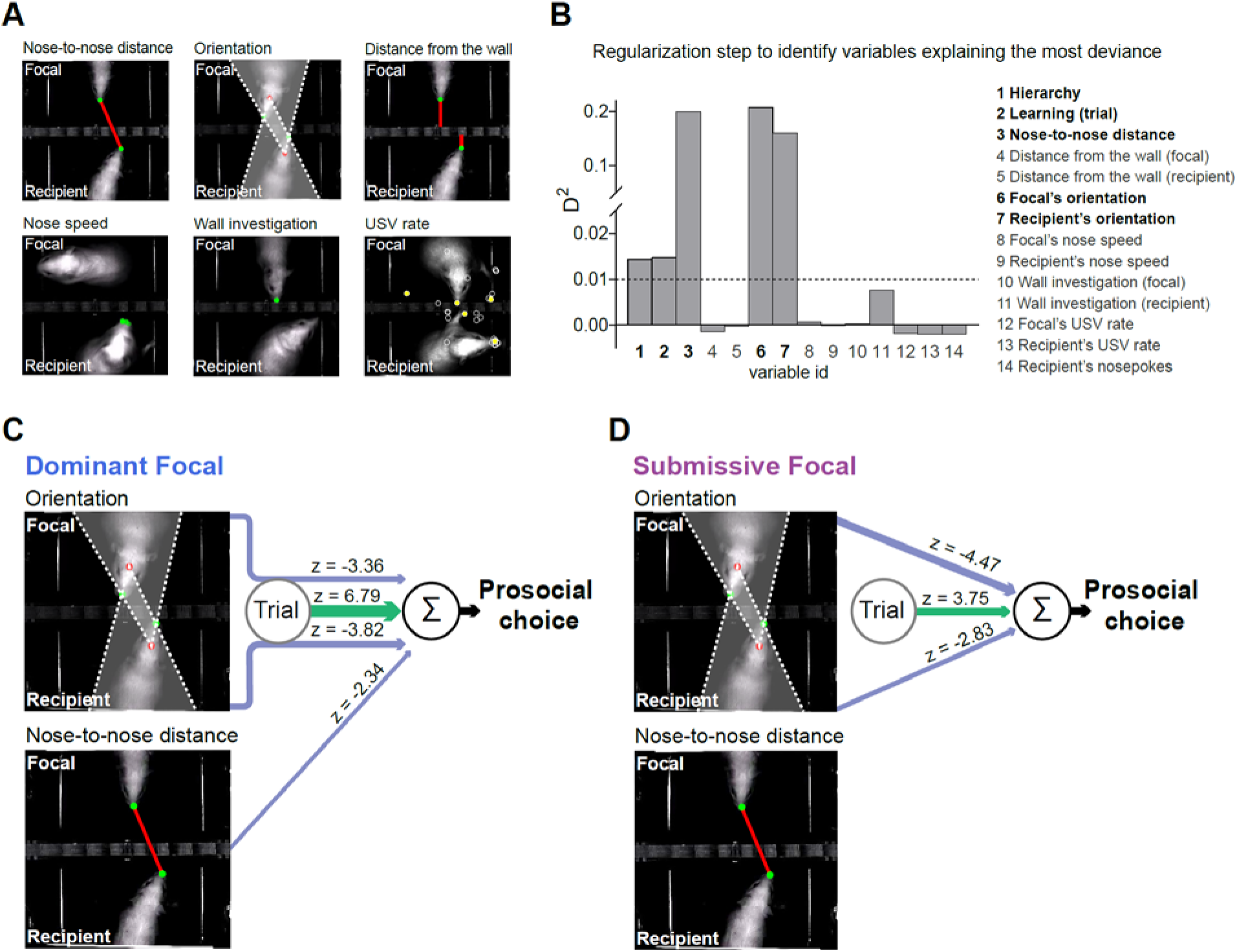
Behavioural predictors of prosocial choice on a trial-by-trial basis. To examine the contribution of a multitude of behavioural parameters to the focals’ choices, we employed a multi-step Generalised Linear Model (GLM) approach. (**A**) Images of the behavioural variables measured either in focal or recipient animals that were included in the analysis, together with the hierarchical status and trial number as a proxy for learning. (**B**) These 14 behavioural and categorical parameters were chosen as regressors and evaluated by their contribution to the explained deviance of the model. The graph on the right shows the mean proportion of deviance (D^2^) computed for each regressor. Trial number, hierarchy, nose-to-nose distance, focal orientation angle, and recipient orientation angle were selected as the regressors that explained more than 1% of the deviance (dashed line) and used to fit a reduced GLM (see **Figure S6** for unique contribution analysis of these variables). Because the latter 3 regressors were found to interact simultaneously with both trial number and hierarchy (**Supplementary Table 2**), we fitted separate GLMs for dominant and submissive animals, thus removing hierarchy from the models and facilitating the interpretation of interaction terms (Supplementary Tables 3 and 4). (**C**) Diagrammatic representation of a trial-by-trial GLM analysis for pairs with dominant and (**D**) submissive focal animals. Here, model terms are represented diagrammatically: The Σ symbol represents the summation of parameters that influence choice, green lines indicate regressors whose contributions correlate positively with prosocial choice, blue lines indicate regressors whose contributions correlate negatively with prosocial choice. Line passing through “Trial” indicate that the interaction between that regressor and trial contributed significantly to choice and line thickness indicates the strength of those contributions as measured by the z score of the regressor weight. The absence of a line from a behavioural parameter to the Σ symbol represents the absence of a statistical contribution to choice for that parameter.

However, this learning was not observed in pairs with submissive decision-makers (**Supplementary Table 4**). Moreover, the nose-to-nose distance was negatively predictive of choice (the lower the distance between the animals the more it predicted prosocial choice), a relationship that was not observed in submissive decision-makers. Interestingly, this relationship of social distance only in pairs with dominant focals was independent of trial progression, indicating that this regressor was a qualitative characteristic inherent to social status evident since the first interactions in the maze (**Supplementary Table 3**).

## DISCUSSION

In this study, we show that rats’ prosocial tendencies are modulated by social hierarchy, with faster emergence and increased prosociality by dominant decision-makers towards submissive recipients. The emergence of prosocial biases depends on the social dynamics established within the dyads. Submissive animals, when in need of help, are influenced by the behaviour of their dominant decision-makers, follow them more and display multimodal cues that facilitate social synchrony and accelerate learning about choice impact on others by dominant rats. On the other hand, when dominant animals are the ones in need of help and not in control of the situation, they try to obtain their own food displaying clear food seeking behaviour, but without approaching and orienting towards their submissive focal animals, which are the ones making the decisions.

In nature, social hierarchy has an important role in social organisation, survival, reproductive success and health of animals in a group^47, 48^. Rats are very social animals that live in large communities in the wild and display rich social interactions^47, 49^. Our findings show how dominance status modulates prosocial choice in a dyad with established social hierarchy. Prosocial modulation “down the hierarchy” has been also observed in some species of non-human primates^12, 25, 29, 50, 51^ (see Cronin 2012 for review^52^), indicating that the effect of dominance status on prosocial behaviour may be conserved across species. Previous work described that high-ranking long-tailed macaques, *Macaca fascicularis*, are more prosocial than low-ranking macaques, hypothesising that prosocial behaviour is not used by subordinates to obtain benefits from dominants, but by dominants to emphasise their dominant position^12^. Furthermore, the relative dominance position mattered more than familiarity between the partners in modulating prosocial behaviour in this species^29^. Other studies in macaques^25, 50^ and capuchin monkeys^51^ also reported prosociality directed down the hierarchy, while in chimpanzees the direction of prosociality remains less clear^9, 53^ (but see^10^). Less effort has been devoted to the detailed investigation of the behaviours explaining these effects. We addressed this point, analysing how dominance status affected different behaviours of both rats during decision-making. Importantly, we found that submissive recipients were the ones that adapted the most their behaviour towards the dominant focals before their choice; they showed closer proximity in social distance, more direct orientation towards the focal, and their 50-kHz call rate correlated with focals’ prosociality. The cues displayed by recipients were multimodal, and analysis of the predictive value of this information to prosocial choices on a trial-by-trial basis pointed towards body language as the learning signal that animals used to drive their prosocial choices. Indeed, the position of submissive recipients affected the behaviour of their dominants, whereas this influence was not observed in couples where the recipient was the dominant of the pair. We observed an increased bidirectional influence in couples with dominant decision-makers, compatible with increased body movement coordination. Dominant recipients mainly directed their attention to the access to the food, whereas submissive recipients also directed their attention to the focal animal when it was going to choose the selfish option. We propose that these multimodal cues displayed by submissive recipient rats may enhance the social salience of signalling need, when the probability to receive help is low, facilitating the emergence of prosocial choices by dominant rats over testing. Our findings are consonant with studies of helping behaviour in chimpanzees, showing that recipients’ communicative displays towards the donor (begging direct requests) increased the success of receiving help^10, 53^.

Furthermore, our results showing increased proximity in the more prosocial pairs are consistent with a study in humans demonstrating that cooperation in a prisoner’s dilemma game decreased with the physical distance between the players^54^. Moreover, both humans and monkeys have been found to shape social attention according to their relative social status, with individuals preferentially allocating attentional resources to high-status conspecifics^55–57^. This mechanism is likely to be particularly relevant for low-status individuals who more heavily depend on high-status individuals and may allow them to monitor and attend more closely the behaviour of their leaders. In the context of prosocial decision-making, this helps to explain why recipient rats direct more attention to a conspecific controlling and deciding on their reward when this conspecific is dominant. Future studies should investigate whether these higher levels of prosociality displayed by dominant animals are a consistent trait that can be observed when interacting with novel submissive rats from other cages, or whether it emerges through the interactions of an already established hierarchy with a cage mate. In the same direction, it would be interesting to study whether submissive animals are better at displaying need in different contexts, not only in reward-related tasks, i.e., when in need of help to avoid a danger. Moreover, although the identification of stable social hierarchies in females has been elusive until now, it would be interesting to investigate whether prosociality is modulated in a similar manner in female social hierarchies.

We did not find evidence of an effect of familiarity or sex of the interacting animals in the emergence of prosociality. Our results support recent reports that these factors do not modulate emotional contagion nor rats’ prosocial tendencies towards animals under stress^19, 22, 58, 59^ (but see^60^) and extend them to appetitive reward-related contexts, which have been less studied. Previous works have shown that different degrees of familiarity do have an impact in the levels of prosocial behaviour under stress, where the effect of familiarity is not observed when comparing cage-mates versus non-cage-mates of the same strain, but it does when comparing familiar and unfamiliar strains. Future studies should address whether strain familiarity might affect prosocial choices in our reward-based task.

Through detailed quantification of social interactions, we describe the importance of social dynamics and information flow as factors underlying the effects of hierarchy on social decision-making, highlighting the role of the recipient and pinpointing multimodal cues in social distance as behavioural salient correlates motivating prosocial behaviour. Our work has identified complex behavioural dynamics that emerge during social decision-making, paving the way for the study of the neural circuits by which the brain monitors others’ actions to guide social decisions, a complex process which is dramatically affected in several psychopathologies, such as autism spectrum disorders^61, 62^. Importantly, these social dynamics were only observed when social interactions were studied over distance and included into the equation the behaviours not only of the decision-maker but also those of the recipient of help. Strikingly, standard measures of social interaction were blind to the rich social dynamics that emerged in this social decision-making task. This highlights the importance of embracing the complexity of social interactions by expanding their analysis to more quantitative and sophisticated venues and taking into account the contribution of each individual to the joint social interaction. This is still scarce in mechanistic studies regarding the neural circuits of social behaviour, with some recent notable exceptions^23, 59, 63–66^, and invite us to adopt a more complex approach in the study of social behaviour going beyond the standard measurement of direct social investigation of one of the interacting individuals.

## ACKNOWLEDGEMENTS

This work was supported by grants of the NARSAD Young Investigator Grant from the Brain & Behavior Research Foundation under the grant number 26478 to C.M., and the Spanish Agency of Research (grant RTI2018-097843-B-100 to C.M.). M.J.M.G. was supported from “la Caixa” Foundation (ID 100010434), under the agreement LCF/BQ/DI17/11620002 and from Fondazione Ing. Aldo Gini. D.A.L. received a travel grant from the Carolina Foundation in the context of this collaboration. We acknowledge Diana F. Costa, member of our team, for drawing illustrations used for figure 1. We thank Dr. Daniel Takahashi (Brain Institute, Federal University of Rio Grande do Norte, Natal, Brazil) for helpful discussion on the use of Partial Directed Coherence and Dr. Jaime de la Rocha (Institut dʹInvestigacions Biomèdiques August Pi i Sunyer (IDIBAPS), Spain) for his help in planning and discussing the GLM analysis. We thank Victor Rodríguez Milán from the Scientific HARdware and Electronics service (SHARE) of the Instituto de Neurociencias, for his precious help in assisting with the customization and optimization of the behavioural setup and the behavioural facility, Dr. Gonçalo Lopes (NeuroGEARS Ltd) and Dr. Aarón Cuevas (Universitat Politècnica de València, Spain), for their help in implementing data acquisition and Mª del Mar Francés Pérez for her support with animal care and maintenance. We thank Prof. Carmen Sandi (Brain Mind Institute, École Polytechnique Fédérale de Lausanne (EPFL), Switzerland) for invaluable comments and feedback on the manuscript.

## AUTHORS CONTRIBUTION

Conceptualization, C.M. and M.J.M.G; methodology, software, formal analysis, M.J.M.G., J.E.A., K.C., and D.A.L.; investigation, M.J.M.G., A.S.M., and H.B.G.; resources: D.A.L., writing – original draft, M.J.M.G. and C.M.; writing – review and editing, M.J.M.G., C.M., D.A.L., K.C, J.E.A.; visualisation, M.J.M.G., D.A.L. and J.E.A; supervision, C.M.; funding acquisition, C.M.

## STAR METHODS

## RESOURCE AVAILABILITY

### Lead contact

Further information and requests for resources should be directed to and will be fulfilled by the lead contact, Cristina Márquez

### Materials availability

This study did not generate new unique reagents.

### Data and code availability

● All data reported in this paper have been deposited at Mendeley Data and are publicly available as of the date of publication. DOIs are listed in the key resources table.
● Original codes for Granger causality analysis and multi-step GLM have been deposited at Mendeley Data and are publicly available as of the date of publication. DOIs are listed in the key resources table.
● Any additional information required to reanalyse the data reported in this paper is available from the lead contact upon reasonable request.

## EXPERIMENTAL MODELS AND SUBJECT DETAILS

### Subjects

86 adult Sprague-Dawley rats, 74 males and 12 females (OFA, Charles-River, France) were used in this experiment, being 8 weeks old and weighing between 226-250 g upon arrival to our facilities. Rats were pair-housed and maintained with ad libitum access to food and water under a reversed light cycle (12 hours dark/light cycle; lights off at 8:30 am) in controlled temperature conditions, and with a transparent red tunnel as environmental enrichment (8 cm diameter, Bio-Serv, # K3325). Rats were left undisturbed in their home-cages for two weeks, except for maintenance routines, allowing them to acclimatise to our Vivarium Facility, to reverse their circadian rhythm and start establishing their social hierarchy. After this period, animals were handled six times during two weeks, allowing them to habituate to the experimenter and to eat the new pellets, which were delivered inside the shavings or from a feeder magazine placed inside the homecage. Rats were 3-3.5 months old when starting the prosocial choice task. Experiments were performed during the dark cycle, waiting at least 1 hour and 30 minutes after the lights were off to start with behavioural procedures. Animals were provided by a commercial company, thus previous social experience, social status and degree of relatedness between the animals was not known. Animal husbandry and all experimental procedures were performed following Spanish Guidelines under the code 2016/VSC/PEA/00193 approved by the Dirección General de Agricultura, Ganadería y Pesca of the Generalitat Valenciana, which are in strict compliance with the European Directive 86/609/EEC of the European Council.

## METHOD DETAILS

### Modified food competition test

After acclimation to the vivarium and handling by the experimenter, 23 pairs of male cage-mates rats were tested in the Food Competition Test (mFC)^30^, to identify the stable social hierarchies. This tests reliably measures already established social hierarchies by introducing a subtle conflict for the access to palatable pellets in the homecage of non food-deprived pairs of animals. Briefly, for this test the homecage lid was replaced by a modified laser-cut acrylic one incorporating a fully transparent feeder for hosting the pellets (Dustless Precision Pellets, 45mg, Rodent Purified Diet). The feeder was designed so only one animal could access the palatable pellets at a time, leading to subtle conflict and competition for the reward. Moreover, the feeder counted with a sliding door to prevent the access to the pellets during inter-trial intervals and an opening on the top where the experimenter could deliver the pellets in each trial. Before testing, rats were habituated for three consecutive days to wait for the sliding door to open and to eat the pellets individually, while the partner rat was kept in a separate cage. In each day of habituation, the fur of the animal was marked to facilitate identification from video and the rat was placed alone in the homecage, with the new lid hosting 10 pellets per trial. In the habituation sessions, the structure of the trial was the following: the rat was allowed to explore in his home cage for 2 minutes with the sliding door of the modified lid closed, thus preventing access to the 10 pellets. Then the feeder was open and the rat had access to the 10 pellets for a period of 2 minutes, after which the sliding door would close for an inter-trial interval of 2 minutes. The number of trials in the habituation ranged from 2 to a maximum of 4 trials per daily session. After habituation, the pairs of cage-mates were re-marked and tested for 2 consecutive days in a social context, inducing now the competition for the positive reinforcers. In the test, a trial started with 1 minute of exploration, with the sliding door of the modified lid closed and hosting 10 pellets. Then the feeder was open and the rats had access to the 10 pellets for a period of 2 minutes, after which the sliding door would close for an inter-trial interval of 1 minute. In each day, rats performed 5 trials, having access to a total 50 pellets in a session of 15 minutes. To control for a possible mere effect of performing the modified Food Competition test on prosociality, we divided the animals into two groups: one group was tested twice for social hierarchy (n = 10 pairs), with two sessions of the modified Food Competition test performed before the start of individual training for the Prosocial Choice Task (PCT), followed by two sessions performed after the PCT, whereas the second group (n = 13 pairs) completed two sessions of the modified Food Competition test only after the being tested PCT (**Figure S1K**). Consumption was quantified by video annotation and the total number of pellets eaten by each animal over the days indicated the social status between them. Statistics evaluating differences between dominant and submissive animals were performed in the average consumption of all days. To have a quantitative measure of the strength of the differences in hierarchy across dyads we computed a Dominance Index (DI), as previously proposed^30^. Briefly, the pellets eaten by the recipient are substracted from those of the focal and normalized by the total number of pellets eaten:

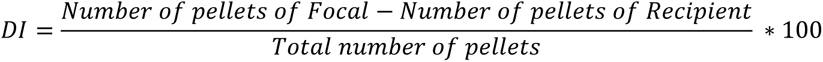

The sign of the DI indicates whether the focal is dominant (positive values) or subordinate (negative values). One pair of animals displayed differences in pellet consumption between the interacting animals smaller than 5%. In this pair social hierarchy was not reliable and categorisation of dominant and submissive was not possible, thus this pair of rats was excluded from the study.

### Prosocial choice task (PCT)

The propensity to perform actions that benefit others was evaluated in the prosocial choice task (PCT), where 43 pairs of non-food deprived rats were tested in a double T-maze, one per animal, as previously described^17^. The two individual mazes (one for the decision maker and the other for the recipient of help rat) are separated by a transparent perforated wall, thus allowing rats to see, hear, smell and partially touch each other. In each maze, a central arm gives access to a choice area and two reward areas where food is delivered in food magazines (**Figure 1A**). Access to the reward area is prevented by automated doors, controlled by nose ports placed above them. Rats had to poke on a nose port for the door underneath to open, thus allowing them to enter the reward area, reach the food magazine and run around the maze back to the choice area, initiating a new trial. For each pair, one rat was assigned to be the focal (decision-maker) and the other the recipient. Rats learned individually to move around the maze and retrieve pellets before the social task. After individual training, rats were tested in the PCT for five consecutive daily sessions of 40 minutes, during which they could perform trials *ad libitum*. A trial would start when both animals were present in the central corridor, giving simultaneous access to the choice area. There, recipient animals could display food-seeking behaviour by performing nose pokes on the side where they would expect the reward. Then, focals could choose between poking on the same side of the recipient, providing access to the lateral arm where both animals would receive one pellet (prosocial choice) or poking on the opposite side, entering the lateral arm where the focal would receive one pellet and the recipient none (selfish choice) (**Figure 1B**). In both choices, focal and recipient rats went to the same side of the maze, and returned to the central corridor to reinitiate a new trial.

Different types of pairs were tested in the PCT. To study the role of **familiarity** as a possible modulator of prosocial choices, two independent groups of male rats were tested: one where decision-maker and recipient were familiar animals (n=13), defined as cagemates living as a stable dyad for at least 1 month before behavioural testing; and another group where decision-maker and recipient were unfamiliar (n=14), defined as rats from the same strain that were not cagemates, that met for the first time in the PCT, and were maintained over the rest of sessions while not being cage mates. Furthermore, we studied the role of the **sex** of the interacting dyads by comparing the prosocial levels of males and females, both of the groups composed of familiar dyads (i.e male cagemates (n=13) were tested together and compared to dyads of female cagemates (n=6)). Finally, the role of **social hierarchy** was evaluated by comparing two independent groups of male cagemates dyads, which lived together for at least one month before behavioural testing. In one group the dominant animal was the decision maker of the pair and would decide whether to provide food to its submissive cage-mate (n=9), while in the other group the submissive animal would decide whether to be prosocial or not to its dominant partner (n=13).

### Behavioural apparatus for the Prosocial Choice Task

The setup consists of two identical, fully automated double T-mazes (Gravaplot, Sintra, Portugal), that are automatically controlled using Graphic State 3.03 software and the Habitest interface (Coulborun Instruments, Allentown, PA, USA). Custom-made automatic doors (WGT-Elektronic, Kolsass, Austria, and Mobiara R&D, Lisbon, Portugal) triggered by infrared beams control the positions of the rats in the mazes, such that when the rats activate the beam a specific door would open, allowing the animals to move to a different area of the maze. Each T-maze has a central corridor as starting point, which gives access to a choice area through an automated door. The choice area is flanked by two lateral reward arms, at the end of which there is a food magazine. To enter the lateral arms, rats had to poke in a light-cued nose port to activate the infrared beam controlling the door underneath. The moment when the focal animal pokes in one of these nose ports, thus opening the doors of the corresponding side that give access to the reward area, is defined as the moment of the decision, i.e when the focal animal reports its choice. Once in the lateral arm, rats could retrieve the food (one pellet per trial), triggering the opening of the door that gives access to a small runway leading to the starting point at the central corridor, thus initiating a new trial. Before being tested in the PCT, rats were trained individually. The roof of each maze consisted in transparent, 2 mm-thick acrylic walls, being perforated to facilitate the detection of ultrasonic vocalisations by the microphones above them. In addition, a transparent, 2 mm-thick acrylic wall was positioned on top of the central wall separating the two mazes and between the microphones to facilitate call assignment. During individual training opaque acrylic walls were placed in each T-maze, thus isolating them, covering the communicating holes and preventing the rat in one maze from seeing the other maze. After the individual training, the opaque acrylic walls were removed and the PCT started.

### Individual training

All animals were habituated to the maze environment for 4 daily sessions of 15-20 min each. Rats were allowed to explore the maze and retrieve the pellets that the experimenter previously placed over the floor of the maze and in the food magazines. In addition, the doors of the maze were manually activated so that the animals could habituate to the noise produced by their opening/closing. After habituation, individual training started. On the first day, all animals were shaped to rear to poke in the nose port for opening the door that gave access to the food magazine. Rats could enter both arms that were rewarded with one pellet per trial. After this first day, each rat of a pair was randomly assigned to be the decision-maker (focal) or the recipient. From this moment, focal and recipient rats received distinct kinds of individual training, for a maximum of 12 daily sessions of 20-30 minutes each. Focals learned to perform one poke on any side of the choice area, to access the lateral arms in order to retrieve the pellet and go back to the central arm to start a new trial, until they reached a performance of at least 1.5 trials/minute. Rats tend to alternate, and no side preference was observed at the end of the training (baseline). For recipients, only the nose port on the rewarded side was active. Thus, recipients learned to poke only to one side, and the number of nose pokes required to open the automated door gradually increased over training, to ensure food-seeking behaviour and clear side preference (for further details on nose poke training, see Marquez et al., 2015). In the last 4 sessions, after nosepoking on the preferred side, the opposite door would open and recipients were forced to visit the unrewarded arm in 10 and 20% of the trials. In this manner, recipients would learn that even if no pellet was delivered in the unrewarded side, they would have to enter that lateral arm and go back to the central corridor to start a new trial. Finally, recipients were briefly re-trained immediately before each session of the PCT, to prevent extinction of food-seeking behaviour. Focal and recipient role were fixed throughout the entire experiment.

### Video and Sound acquisition

All the experiments were performed during the dark phase of the animal’s light cycle and video recordings were captured at 30 frames per second and 1280 x 960 pixel resolution under infra-red illumination (PointGrey Flea3-U3-13S2M CS, Canada, FlyCapture). We used two cameras, each positioned above one double maze, and centred on top of the choice area. Ultrasound was recorded at a sampling rate of 214285 Hz with two externally polarised condenser microphones (CM16/CMPA) connected to an UltraSoundGate 416H (Avisoft Bioacustics). We positioned two microphones on top of the choice area of each double maze, one microphone per T-maze (44 cm from the floor of the maze and 15 cm from the acrylic wall between them). For each session, video and audio acquisition start was simultaneously triggered through a common TTL delivered from visual reactive programming software Bonsai^37^ through an Arduino Uno (ARDUINO®). With Bonsai, we also sliced the entire session videos into video chunks, corresponding to the choice period of each trial. For the synchronisation of the video with the data obtained from the interface controlling the mazes, we extracted the timestamps from the coulbourn interface, and tracked the blinking of an infra-red LEDs placed in the visual field of the camera which was triggered at the time of each start trial and focal’s choice. Sound recordings were synchronised to video data aligning each start trial with the recorded sound of the opening of the door that gave access to the choice area.

### Pose estimation of unmarked socially interacting animals via Bonsai-DeepLabCut

A custom workflow of the Bonsai-DLC interface^67^ (Python 3, DLC, version 2.2) was used to track unmarked body parts of the animals on both sides of the double maze simultaneously. Tracking was performed offline for all single trials’ videos from the first two sessions of the PCT, with a confidence threshold set to 0.7. Video analysis’ temporal resolution was determined by the camera’s acquisition frame rate (33 ms), whereas spatial resolution was calculated by measuring a reference known distance in pixel values at the height where animals move in the maze (spatial resolution 0.59 mm). A cropped image corresponding to the choice area of one T-maze was used as region of interest (ROI) to train the model, applying an offset for each of the choice areas to maintain the original frame coordinates. DLC was trained on videos with one animal in the T-maze (26 videos from different animals). 25 frames per video (650 frames in total) were annotated and used to train a ResNet-50 neural network for 600,000 iterations.

For each video frame in the recordings, we obtained the location of the noses used to compute the euclidean distance between the rats in the choice area, and the y coordinate of the nose to retrieve the distance of each animal from the central wall. Moreover, we calculated the position of the nose of each animal in the maze, where movements in the x coordinate would indicate movement towards the selfish or prosocial port (being coordinate 0 cm the position of the selfish side and 17 cm the position of the prosocial side). In our task, focal and recipient animals of different pairs are counterbalanced when assigned to a side of the double maze, such that in some pairs the focal rat would appear in the upper part of the video and the recipient rat on the bottom part (as illustrated in **Figure 1B**), while the opposite occurs in the rest of the pairs. Furthermore, the prosocial side is also counterbalanced, such that it would be to the right side for some focals and to the left side for the remaining ones. Thus, we moved and scaled tracking data from different recordings to a common reference space. For this, we used the coordinates of the central wall of the double maze as space scale factor for pixel to meter conversion.

Orientation of one rat towards the other was computed as the angle between the vector from the middle of its head (halfway between the ears) to its own nose and the vector from the middle of its head to the other rat’s nose. We obtained the nose instantaneous speed from the rate of change in its position. For this, we smoothed the nose position time series by independently convolving its two coordinates with a Gaussian window of 0.25 s (full width at half maximum). For each time point, we obtained the velocity vector as the derivative of each smoothed coordinate and computed instantaneous speed as its norm.

### Automatic detection and assignment of Ultrasonic Vocalisations (USVs) in the Prosocial Choice Task

We automatically detected and assigned USVs as thoroughly described in Sirotin et al, 2014^42^. Briefly, USVs were detected from the raw sound recordings with custom built MATLAB routines (The Mathworks). We first obtained the sonograms for each microphone, with a 0.25 ms time step and detected times with low entropy (<6.5 bits) of the frequency spectrum in the 18-100 kHz range. We then defined as USVs segments of low entropy, those lasting at least 3 ms and bounded by silence of >20 ms. USVs were then curated by automatically discarding as noise those with high power in the sonic range (5-18 kHz) and visually inspecting the sonograms, removing any noises detected as USVs by mistake. Next, each vocalisation was assigned to either the focal or the recipient rat, by comparing the signal from both microphones. USVs that crossed the entropy threshold in only one microphone were assigned to the rat on the T-maze below it. If the same USV was picked up by both microphones, we assigned it to the rat under the microphone with lowest entropy values. Rats vocalising at the same time will typically produce USVs with non-overlapping fundamental frequencies. When simultaneous signals from both microphones were found to differ by at least 1kHz during >3 ms, we concluded that both rats vocalised simultaneously and assigned to each one the USV detected by the microphone on its side. As in Sirotin et al., 2014^42^, we used recordings with only one rat in the double maze to validate the USV assignment, yielding an accuracy of 94% (**Figure S4**).

### Multimodal analysis of USVs and tracking data

We temporally aligned audio and video of each recording session. This allowed to retrieve the video time and frame when a USV was emitted, tagging each USV with relevant behavioural information, i.e.- in which trial the USV was emitted and the location of the noses at the time a USV was emitted during the choice period (**Figure 5B**). We were then able to selectively quantify the USV number and rate during the interaction time (noses of both rats simultaneously detected in the choice area), which we used for **Figures 5, S4C and S5**. After moving and scaling tracking data to a common reference, we were able to map the location of the noses at the time of USV emission from all recording sessions (**Figure 4C**).

### USVs Classification

We performed automated classification of the USVs that were already detected and assigned to the emitter rat from the dominance groups (n=45.898) into three different classes of 50kHz USVs that correspond to different vocal programs: “flat”, “frequency-modulated” and “step”. We extracted a grayscale image of the sonogram for each vocalisation (sonogram duration 200 ms, frequencies 25-100 kHz) and used a convolutional neural network for supervised image-based classification following the pipeline in VocalMat^45^. We manually selected and labelled flat (n=1002), frequency-modulated (n=1003) and step (n=921) calls, equally distributed across animals, randomly assigned 90% of each class as training set and trained the network using the original weights from VocalMat as starting point (original script available here: https://github.com/ahof1704/VocalMat/blob/master/vocalmat_classifier/training/train_model.m). Classification accuracy measured on the test set was 98%.

## QUANTIFICATION AND STATISTICAL ANALYSIS

Data obtained from the interface controlling the mazes, video analysis and USVs recordings was parsed and processed with Python (Python Software Foundation, version 3, https://www.python.org/) and MATLAB (version 2018a, The Mathworks, https://matlab.mathworks.com/). Probabilistic statistical analyses were performed using IBM SPSS Statistics version 23 for Windows (https://www.ibm.com/analytics/spss-statistics-software) and Bayesian statistics with JASP version 0.16.1 (https://jasp-stats.org/). GLM analysis was performed using the package modEvA^68^ from the R Project for Statistical Computing (https://www.R-project.org/). Normality of the data was tested with the Shapiro-Wilk method.

### Analysis of social interactions prior to choice

All the analysis of social interactions prior to choice was restricted to the video frames where both focal’s and recipient’s noses were tracked in the choice area (the ROI for the training of DLC). We called this portion of the total choice time “**interaction time**”.

#### Duration of mutual direct investigation

The duration of mutual investigation in a trial was calculated from the frames in which the nose-to-nose distance was shorter than 2 cm. Absolute thickness of central wall of the double maze was 1 cm, however, after manual observation of the nose coordinates we decided to expand the distance to 2 cm in order to include in this measure all the mutual investigations that would happen in a diagonal, mostly across two separate holes in the perforated wall.

#### Wall investigation of each animal

as a proxy of time sniffing through the wall, rat’s wall investigation was calculated from the frames in which the distance between the rat’s nose and the central wall was equal to zero (**Figure 3M** and **S3N-O**).

#### Quantification of social interactions in social distance

For each trial, we extracted the median of the time series of the different variables (the nose-to-nose distance, nose distance from the central wall, nose speed and orientation towards the partner). We then averaged the medians from all trials of a pair/rat to obtain a value for each subject, that we used for statistical analysis.

#### Radar plots of head orientation towards the partner

For the radar charts in **Figures 3I, 3K, 3L** and **S3M** we retrieved in each trial all the frames with orientation value, which ranged from 0 (rat oriented to the nose of the partner) to 180 degrees (rat oriented to the opposite direction of the nose of the partner). We then calculated the percentages of frames belonging to each of three ranges (0-60 indicative of more direct gazing, 60-120 and finally 120-180 indicative of positions where one animal oriented opposite to the partner). For “overlapping orientations”, we calculated the percentages of frames in which the orientations of both rats fell within the same range.

#### Visualisation of early dynamics of social interactions in social distance

We selected, for each trial, the frames (time points) where both noses were tracked (interaction time). Next, we aligned the new time series, so that the first frame in each series was set as time 0. We then obtained an average time series for each hierarchy-trial type condition, by averaging the time series of the different trials at each interaction time point, up to the median duration of the interaction per trial (3.3 sec, line graphs of behavioural dynamics in **Figure 3** and **Figure S3**). Finally, for the statistical comparisons we calculated the median of each new time series corresponding to a trial and averaged the medians of all the trials belonging to the same hierarchy-trial type category (dot graphs with SNK test in **Figure 3** and **Figure S3**).

### Granger causality from Partial Directed Coherence

To assess whether the behaviour of one rat influences that of the other within trials (**Figures 4 and 5**), we applied partial directed coherence (PDC), a frequency decomposition of Granger causality^69^, using routines from the AsympPDC implementation^70^ (package available at http://www.lcs.poli.usp.br/~baccala/pdc/CRCBrainConnectivity). Briefly, for each condition (e.g. “focal animal is dominant”) we fitted a single vector autoregressive (VAR) model to the time series of interest from focal and recipient rats and computed from it the information PDC (iPDC)^71^ spectra from the focal to the recipient and vice versa. We integrated each iPDC spectra to obtain information flow (*I*_flow_), a scalar value representing the causality from one rat group (focal or recipient) to the other in units of information transfer (bits). We used trial-shuffle surrogates and resampling statistics to test for significance of each *I*_flow_ and of *I*_flow_ differences across conditions.

#### Granger causality between focal and recipient positions

In detail, to quantify within-trial causality between the positions of the rats along the x axis (running parallel to the wall separating the rats, from one nose port to the other, scaled such that the prosocial nose port is always represented at the “right”, **Figure 4A**), we began by extracting from each trial the longest uninterrupted interaction between the rats from the early interaction time (the first 3.3 seconds selected for the analysis of behavioural dynamics). Trials not containing an uninterrupted interaction of at least 1 second were discarded from the analysis (93 out of 1998 total trials discarded). We subsampled the data by averaging every 3 time points, resulting in a sampling rate of 10 samples per second. At this point, each trial is represented by a vectorial time series of two dimensions (*x_focal_*(*t*), *x_recipient_*(*t*)) and 10 to 33 time points (1 to 3.3 seconds in duration). We sorted the trials into groups representing each condition and normalized *x_focal_* and *x_recipient_* by subtracting the mean of each variable in the whole condition and dividing by its standard deviation (note we did not normalize the data trial by trial). Next, we fitted a VAR model of order 2 to each individual trial vectorial time series. Our method requires a fixed order for the VAR models and 2 was the median optimal model order for individual trials as per Akaike’s information criterion. We then computed the mean of all VAR models, thus producing a mean autoregressive model for each condition from which we calculated the iPDC spectra from focal to recipient and recipient to focal and we integrated each iPDC across all frequencies into *I*_flow_ as (adapted from equation 8 in^71^):

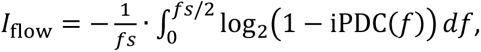

where *f s* is the sampling rate.

We implemented surrogates and resampling statistics to test for significance of *I*_flow_ and *I*_flow_ differences between conditions. We began by performing trial-shuffle surrogates within each condition. To construct each surrogate, we paired the data from the focal in each trial with data from the recipient in a random trial from the same condition. Since trials were of variable duration, we randomly matched each trial only with others having at least its duration and kept only data up to their common duration (481 of 1832 trials were of maximum duration, ensuring well-varied surrogates for all). In this way, surrogate datasets represent the null hypothesis whereby there is no interaction between the two rats within each trial. We obtained the iPDC and *I*_flow_ from each of 1000 surrogates and calculated a one-sided p-value with finite-bias correction^72^ as:

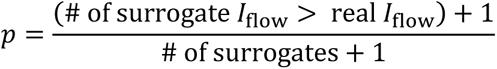

To account for the positive bias in PDC, we subtracted the median surrogate iPDC from the real iPDC before calculating each final reported *I*_flow_ value.

We tested for significant differences between two given *I*_flow_ values by obtaining bootstrap distributions of their differences. For a condition with *n* trials, we get each single bootstrap estimate by selecting *n* random trials with replacement and obtaining iPDC, subtracting the median surrogate iPDC from it and calculating *I*_flow_ as before. We do 1000 subtractions of bootstrap estimates from each *I*_flow_ and compute a 2-sided finite-bias-corrected p-value against the null hypothesis of there being no difference as:

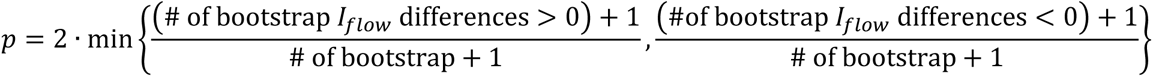

#### Granger causality between recipient USVs and focal orientation towards the emitter

For analysing causality from emission of USVs to orientation of the listener rat towards the emitter (Figure 4F) we followed the pipeline described above for rat positions, with adaptations as follows. We first constructed a binary time series with one sample per video frame valued 1 if the rat emitted a USV with onset in the time interval between that and the next frame and 0 otherwise. We then convolved this with a gaussian kernel of full-width at half-maximum of 0.25 s to obtain a continuous representation of vocal production and added to these time series gaussian noise with a sigma of 10% of their standard deviation as a necessary stochastic component as suggested in ^70^. We then extracted the recipient USV and focal orientation time series from the rats, keeping the longest uninterrupted interaction between the rats for each trial up to 10 seconds of interaction time (1 second minimum duration), downsampled to 10 Hz and proceeded to obtain iPDC and *I*_flow_ from USV_recipient_ to orientation_focal._

### Generalised Linear Model (GLM) Analysis

To examine the trial-by-trial contribution of a multitude of parameters to the focal’s choice, we employed a multi-step GLM approach. First, we fitted a binomial GLM with 14 behavioural and categorical parameters (see **Figure 6**) for regressor details) using the formula

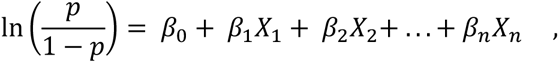

where *p* is the probability of a prosocial choice, *β*_0…n_ are the regressor weights, and *X*_1…n_ are the regressor values. Because Wald tests for statistical significance provide an incomplete interpretation of regressor contributions, we instead employed an alternative approach similar to that previously described by Musall and colleagues^73^, in which we computed the proportion of deviance (D^2^) explained individually by each regressor. To achieve this, for each regressor in the model, we shuffled the values of every *other* regressor’s values, resulting in a dataset in which one regressor contained the actual values on each trial, but all the other regressors’ values were shuffled. Then, we fitted a GLM to the shuffled trial-by-trial data and computed the D^2^. To provide robustness against potential random imbalances in any single shuffling, we repeated this procedure 1,000 times per regressor and took the mean D^2^ value for each one. Thus, we were able to determine the maximum explained deviance for each regressor. Because each regressor’s D^2^ value is computed independently, it does not describe the amount of unique information each regressor contributes to the model. Two regressors could have a similar D^2^, but if they are related or dependent on each other, then their unique contributions to the predictive power of the model will be limited. Therefore, we also computed the ΔD^2^ for each regressor: the proportion of the deviance that is *uniquely* explained by each regressor. To achieve this, for each regressor, we shuffled the values of only that regressor, leaving all others intact. We then fitted a GLM to this dataset and computed its D^2^. Next, we subtracted this value from the D^2^ of the full GLM with all regressors intact to obtain the ΔD^2^. To provide robustness against any single shuffling of the data, we repeated this shuffling procedure 1,000 times for each regressor and took the mean D^2^ for each one. In this manner, ΔD^2^ is essentially a measure of how much predictive power the model loses when each variable is shuffled, thus revealing its unique contribution.

Next, to more deeply examine the regressors that explained most of the deviance as well as how they might interact with hierarchy and trial progression, we fitted a reduced GLM using only those regressors that explained more than 1% of the deviance: trial number, hierarchy, nose-to-nose distance, focal orientation angle, and recipient orientation angle. This time, we also included interaction terms between trial and the remaining variables, as well as for hierarchy and the remaining variables, since our previous observations strongly suggested that other variables may interact with those two.

Finally, to tease apart the resulting triple interaction involving the behavioural variables plus both trial and hierarchy, we fitted separate reduced GLMs on dominant focals and submissive focals, this time using 4 variables: trial, nose-to-nose distance, focal orientation angle, and recipient orientation angle, as well as an interaction term for trial. By fitting separate GLMs to each hierarchical group, we were able to remove hierarchy from the model, thus facilitating a more direct interpretation of the interactions between trial number and the behavioural variables.

### Statistics

Repeated measure (RM) ANOVA with one between-subjects factor and “session” as within-subjects factor was performed to compare prosocial choices between the groups under study (dominant vs submissive focals, familiar males vs unfamiliar males, familiar females vs familiar males, effect of testing for social hierarchy with the modified food Competition test on the prosocial choice task) over the course of the testing sessions. Independent sample t test was performed to assess differences between the groups when examining prosocial choices in each testing day and the average prosocial preference over the 5 days. Paired-sample t test was used for each focal to compare the prosocial choice in each testing day against rat’s baseline preference in the last two days of individual training. One-sample t test was computed for each focal to compare its baseline preference against chance level (50% preference). Bayesian statistics complemented these analyses (Bayesian repeated measures and t test analyses) in order to provide estimates of the strength of the effects. We provide the BF_incl_, BF_+0_ (one-tailed) or BF_10_ (two-tailed) accordingly.

#### Prosocial Choice Index

We computed a prosocial choice index (PCI) to quantify individual differences on choice preference against chance over testing sessions,

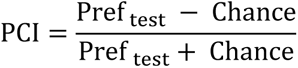

where Pref _test_ corresponds to the proportion of prosocial choices during social testing sessions, and Chance is understood as the proportion of choices equal to 50%. The PCI values show the strength of change in prosocial preference from 50% preference for each rat; [+] PCI show an increase on prosocial preference on social testing sessions compared to chance, [-] PCI show a decrease on prosocial preference from 50%. Distribution of PCIs for each group can be seen in **Figure S1**.

#### Permutation test analysis

To address individual variability on prosocial preference, we performed a permutation test to identify those rats that showed significant change on choice preference against chance. For each animal separately, we generated a distribution of 10.000 permuted PCIs by shuffling the sequences of all choices during social testing with same-length sequences of choices with prosocial preference equal to 50%. Rats then were assigned to three different categories by comparing their actual PCI to the 95% confidence interval (CI) of the distribution of randomized indexes (rat with actual PCI in 2,5% upper bound was considered as *prosocial*, rat with PCI in 2,5% lower bound was considered *selfish*, and those rats with PCI falling inside the 95% were considered as *unbiased*). Lower and upper bound for each individual’s distribution can be found in (**Supplementary Table 1**).

#### Proportions of prosocial, unbiased, and selfish rats

we used χ^2^ test to analyse differences in the proportions of animals classified as prosocial, unbiased, or selfish, for every session of the PCT of the different tested groups (familiar males, unfamiliar males, familiar females, dominant focal males, and submissive focal males).

#### Social interactions

we performed the independent sample t test to assess differences between dominant and submissive focal groups, when examining tracking data extracted and averaged from all the trials of each session. When grouping data by prosocial and selfish trials, A RM-ANOVA with “hierarchy” as between-subjects factor and “choice” as within-subjects factor was used to test for differences between the two groups across trial type. Finally, a one-way ANOVA followed by Student-Newman-Keuls (SNK) post-hoc test was used to evaluate differences among dominance-trial type categories, when examining the early dynamics of the interaction time.

#### Social dominance

dominance index as a measure of the strength of the social hierarchy was correlated with prosociality in the second day using Pearson correlation.

#### Task performance

a RM-ANOVA with “hierarchy” as between-subjects factor and “session” as within-subjects factor was used to compare dominant focal and submissive focal groups in number of trials performed over the 5 days of testing of prosocial choice task.

#### Nosepokes and choice time

difference between hierarchy groups in the average number of recipient’s nosepokes per trial was assessed with the independent sample t-test. For the recipient’s latency to nosepoke, recipient’s nosepoke duration (computed by subtracting the time of the rat’s snout entering the nose port (activating the infrared beam) to the time the rat’s snout exited the nose port (inactivating the infrared beam)) and focal’s choice time, the non-parametric Mann-Witney U test evaluated differences in the distributions.

#### Proportion of USV classes

we used repeated-measures ANOVA to assess differences in the proportions of USVs classified as flat, frequency-modulated, and step, across days of testing and according to the emitter agent (focal, recipient) and their hierarchy (dominant, submissive).

#### Relationship between USV rate and speed

we divided for each rat the number of USVs emitted while its nose was moving within each of four instantaneous speed bins by the total time in each bin (**Figure S4D**). Then, differences in USV rate between focals and recipients was assessed across the different speed bins using RM-ANOVA. For cross-correlations between USV emission and nose speed (**Figure S4E**) we obtained the USV time series as explained for Partial Directed Coherence (without subsampling nor adding noise) together with instantaneous nose speed for each trial. For each rat, we concatenated all its trials leaving gaps of 1 second with missing values between them and run the cross-correlation with a maximum lag of +/- 1 second. For each lag, we normalized the cross-correlation value by the total number of non-missing samples used for its computation to obtain the unbiased cross-correlation estimate. We subtracted from each cross correlation the mean of 1000 within-rat trial-shuffle surrogates.

#### Analysis of partial correlations

We performed partial correlations with nose-speed of the animals as covariate, to study the correlation between USV rate of the different animals and prosociality.

### Video legends

**Movie S1.**
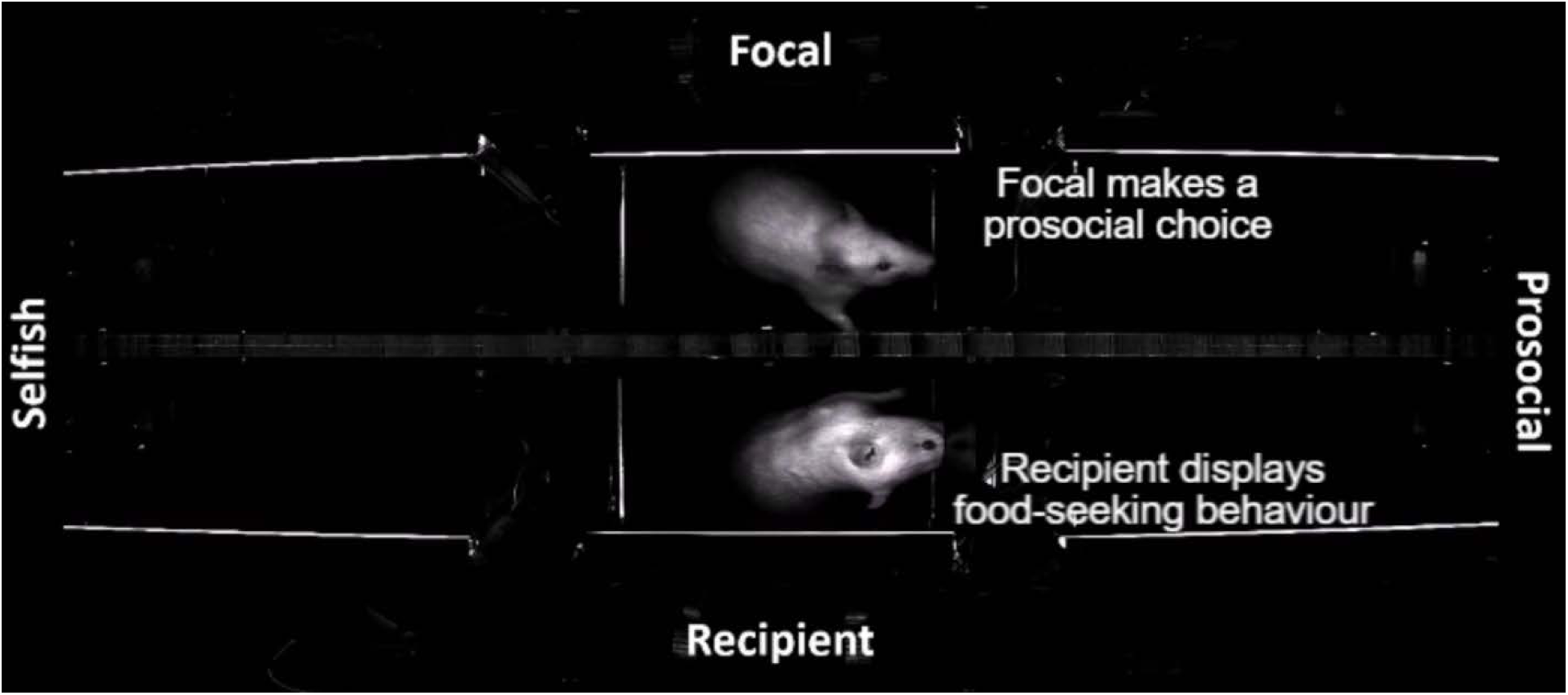
Representative trials of the Prosocial Choice Task. Related to Figure 1A-C. The video shows representative prosocial and selfish trials, from start trial to reward delivery, of a pair of rats performing the Prosocial Choice Task. It can be observed the recipient rat displaying food-seeking behaviour (poking in the nose port on the preferred side), the focal rat making its choices, and how both animals visit the lateral arms where food is delivered.

**Movie S2.**
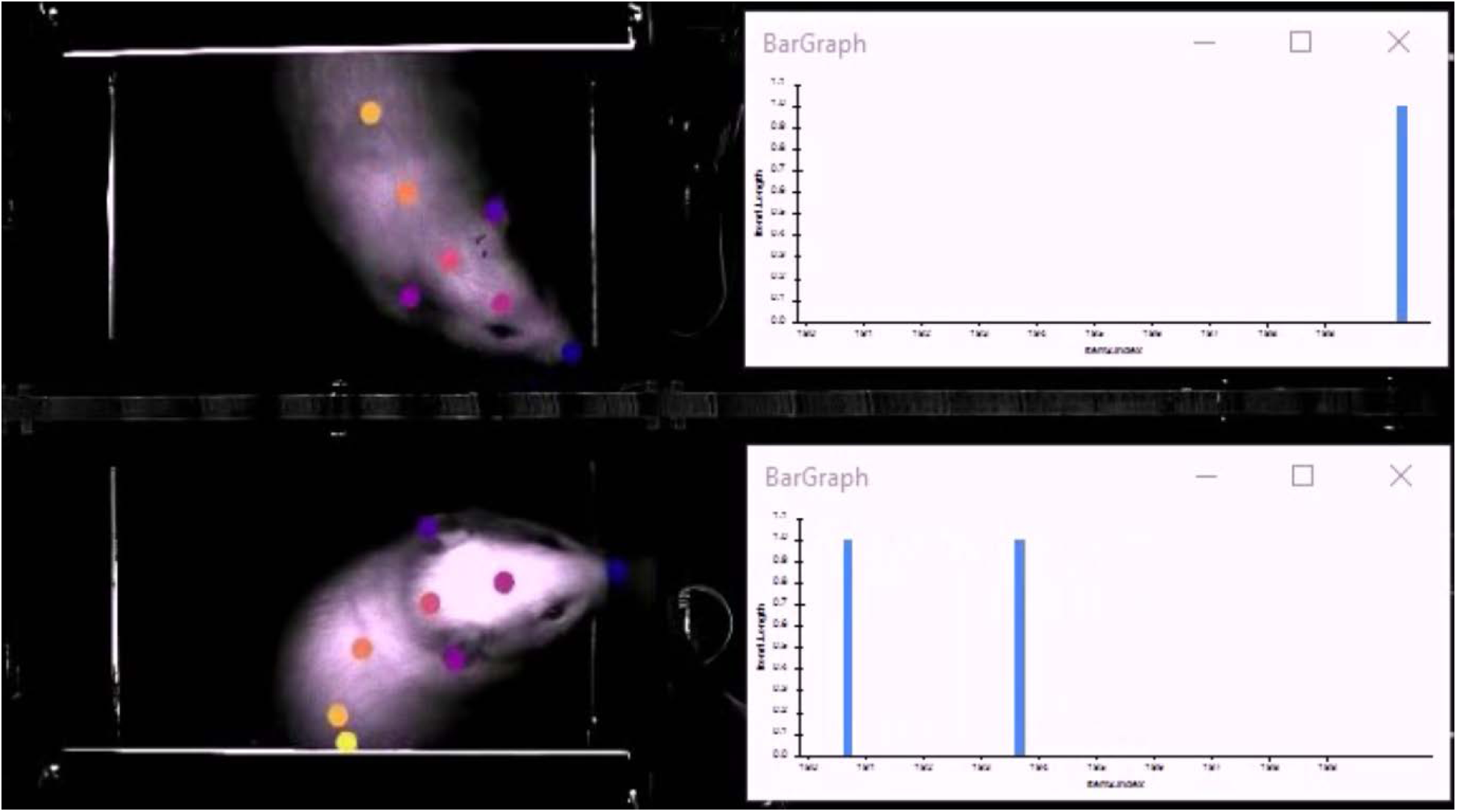
Representative trial with pose estimation and call events. Related to Figure 5. Example video for illustrating tracked animals in the choice area and time of USV emission. Animal body parts are labelled using DeepLabCut. After synchronisation of video and audio data, we built for each rat a binary time-series, where each frame of the video was tagged with 1 if a USV was emitted or with 0 otherwise. We then used Bonsai to visualise the time of USV emission from each animal as train pulses (blue bars), in parallel with the video. To help visualise the video, we assigned different sounds to the USVs emitted by each one of the animals of the pair.

---

**Figure S1.**
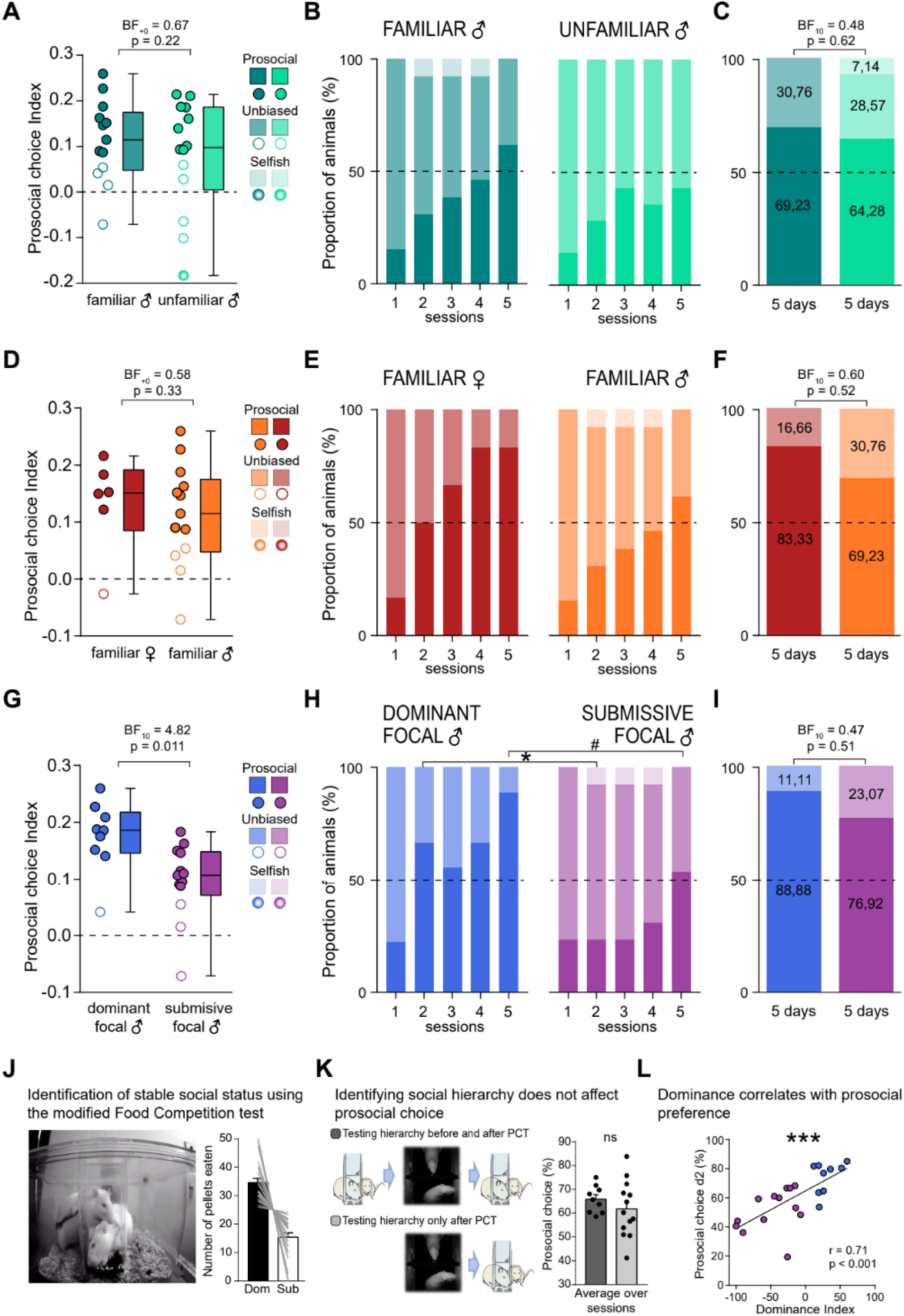
Individual differences in PCT and identification of social dominance. Related to Figure 1. **(A)** Distribution of Prosocial Choice Indexes (PCI, see methods) calculated for familiar and unfamiliar male groups. Each symbol represents the PCI for each rat over the five testing sessions; filled dots indicate prosocial rats, meaning a positive change on preference compared to chance (50%) after a permutation test; empty dots indicate unbiased rats, which preference is not different from chance; and degraded dots represent selfish animals, those having a negative change on preference. Boxplots show median and 1^st^ and 3^rd^ quartiles, wiskers 2.5% and 97.5% percentile values. Independent samples t-test showed no differences between the PCI of the groups t(25)= 0.779, p= 0.222, BF_+0_= 0.67. **_(B)_** We then analysed how the proportion of prosocial, unbiased and selfish animals emerged across testing days. No significant differences were observed between groups, with Bayesian statistics indicating absence of evidence (day1 X^2^ (1, 27) = 0.006, p = 0.936, BF_10_ =0.577; day2 X^2^ (2, 27) = 1.160, p = 0.560, BF_10_ =0.543; day3: X^2^ (2, 27) = 1.122, p = 0.571, BF_10_ =0.492; day4: X^2^ (2, 27) = 1.656, p = 0.437, BF_10_ =0.633; day5: X^2^ (1, 27) = 0.942, p = 0.332, BF_10_ =0.692). **(C)** In the same direction, the proportion of prosocial, unbiased, and selfish rats considering together the five sessions of PCT for familiar and unfamiliar male groups, was not significantly different (X^2^ (2, 27) = 0.964, p = 0.617, BF_10_ = 0.483). **(D)** Same as in A for the distribution of Prosocial Choice Indexes (PCI) for familiar females and males. Independent samples t-test showed no differences between males and females (t(17)= 0.457, p= 0.327, BF_+0_= 0.58). **(E)** Same as B for familiar females and males groups. Although a tendency for female animals to display faster emergence of prosociality could be observed, no significant differences were found across the days (day1 X^2^ (1, 19) = 0.005, p = 0.943, BF_10_ =0.60; day2 X^2^ (2, 19) = 1.644, p = 0.44, BF_10_ =0.67; day3: X^2^ (2, 19) = 1.516, p = 0.469, BF_10_ =0.59; day4: X^2^ (2, 19) = 2.411, p = 0.30, BF_10_ =0.87; day5: X^2^ (1, 19) = 0.903, p = 0.342, BF_10_ =0.71). **(F)** Same as C for familiar females and familiar males groups, where no significant differences between conditions were found (X^2^ (1, 19) = 0.421, p = 0.516, BF_10_ = 0.60). **(G)** Consistent with the percentage of prosocial choices results, PCI where higher for dominant focal males (Independent samples t-test: t(20)= 0.457, p= 0.011, BF_10_= 4.82). **(H)** Dominant focal groups showed significantly different distributions in the second day of testing, indicative of higher number of prosocial animals in early testing days. (day1 X^2^ (1, 22) = 0.002, p = 0.962, BF_10_ =0.54; day2 X^2^ (2, 22) = 6.249, p = 0.044, BF_10_ =5.67; day3: X^2^ (2, 22) = 2.788, p = 0.248, BF_10_ =1.13; day4: X^2^ (2, 22) = 3.046, p = 0.218, BF_10_ =1.25; day5: X^2^ (1, 22) = 3.010, p = 0.083, BF_10_ =1.98). **(I)** However, when taking into account all testing sessions, no significant differences on the proportions were observed anymore (X^2^ (1, 22) = 0.512, p = 0.474, BF_10_ = 0.68). **(J)** (Left) image showing two male cage-mate rats performing the modified Food Competition test for identification of stable social hierarchies in the homecage. In this task, only one of the two animals can gain access to palatable pellets in each trial, leading to a subtle conflict that results in higher consumption of food by one animal of the pair. (Right) Number of pellets eaten by the two rats within each pair averaged across the testing days. The rat eating more pellets over the testing days was categorised as the dominant (‘Dom’) and the rat eating less pellets as the submissive (‘Sub’) of the pair. **(K)** To control for any effect of testing for hierarchy in the modified Food Competition test on prosociality levels, we compared a group of pairs tested before and after the PCT (n = 9), with a group of pairs tested only after the PCT (n = 13). No substantial difference was found between the two groups in the proportion of prosocial choices over sessions, indicating that being tested for hierarchy does not affect prosocial tendencies (independent sample t test: t_(20)_ = 0.989, p = 0.334, BF_10_ = 0.55). Data are represented as group MEAN ± SEM of the prosocial levels of the 5 testing sessions. **(L)** Dominance Index (DI) as a measure of social hierarchy strength positively correlates with prosocial preference displayed during day 2 of PCT testing (r= 0.71, p=0.0002). Blue dots indicate dominant focal rats, and purple dots submissive focal rats. ^#^p<0.1, *p<0.05, ***p<0.001, ns = not significant.

**Figure S2.**
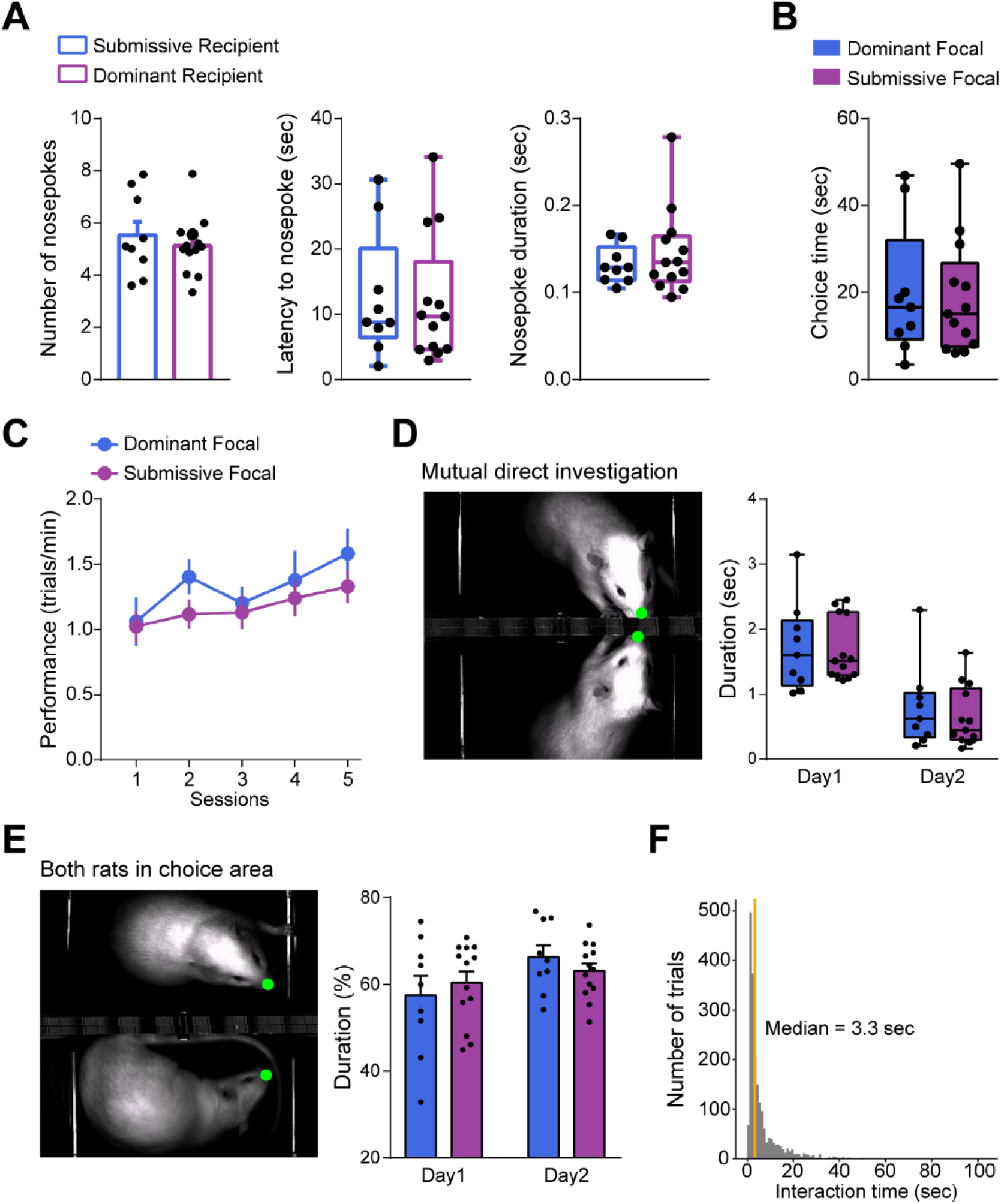
Effects of dominance status on recipient’s nosepokes, choice time, task performance, mutual investigation and interaction time. Related to Figure1, 2 and 3. **(A)** Social hierarchy of the recipients does not affect displays of food seeking behaviour. Submissive and dominant recipients showed no difference in the number of nosepokes per trial (left), nor in the latency to perform the first nosepoke from start trial (middle), neither in the duration of nosepoke (right) prior to choice (independent sample *t* test for the number of nosepokes of submissive against dominant recipients: t_(20)_ = 0.71, p = 0.488, BF_10_ = 0.465; Mann-Whitney U Test for latency to nosepoke and nosepoke duration: *U* = 63, p= 0.794, BF_10_ = 0.384; *U* = 54, p = 0.794, BF_10_ = 0.412). **(B)** The duration of the choice period was similar between dominant and submissive focals (Mann-Whitney U Test: *U* = 61, p = 0.896, BF_10_ = 0.372). **(C)** Dominance status did not affect the performance in the PCT, measured as number of trials per minute (repeated-measure ANOVA with “session” as within-subjects factor and “hierarchy” as between-subjects: “session” (F_(4,80)_=5.577, p=0.004, BF_incl_ = 39.9), “session” by “hierarchy” (F_(4,80)_)=0.695, p=0.535, BF_incl_ = 0.15) and “hierarchy” (F_(1,20)_=0.821, p=0.376, BF_incl_ = 0.61). **(D)** Despite the changes in prosociality observed in day 2, these were not accompanied by differences in the duration of social investigation across the testing days (Mann-Whitney U Test for “mutual direct investigation” on day 1: *U* = 55, p = 0.845, BF_10_=0.415; on day 2: *U* = 65.5, p = 0.647, BF_10_=0.433), nor by differences in **(E)** the percentage of choice time per trial that both rats are present in the choice area, as an index of duration of social interactions in the distance (independent sample t test for “both rats in choice area” on day 1: t_(20)_ = -0.593, p = 0.560, BF10=0.44; on day 2: t_(20)_ = 1.049, p = 0.307, BF_10_=0.57) **(F)** Histogram including all trials from the first two sessions of both dominance groups, showing the median time (orange line) of interaction (total time per trial in which the noses of both rats were simultaneously tracked in the choice area). This value was selected as upper limit for all the trials to visualise the early dynamics of nose-to-nose distance, recipient’s distance from the wall, nose speed and orientation, plotted in Figure 3 and **Figure S3**. Bar graphs in (A) and (E) show MEAN ± SEM and individual values; line plot in (C) shows MEAN ± SEM; box plots in (A), (B) and (D) show median, first and third quartiles, with whiskers indicating maximum and minimum values. Individual values correspond to the mean over days for each animal.

**Figure S3.**
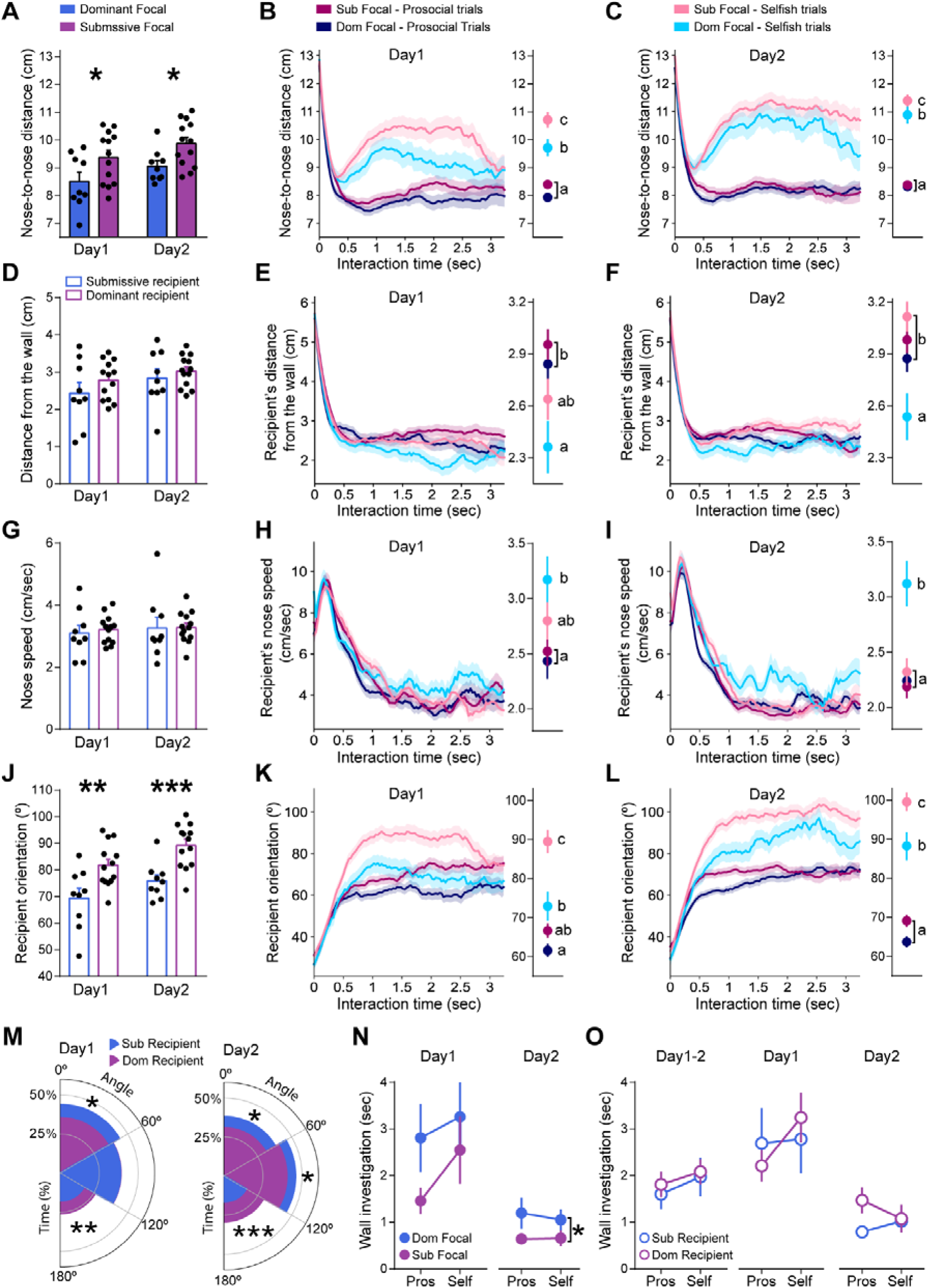
Social dominance modulates the dynamics of social interactions prior to choice across days. Related to Figure3. (**A**) The median distance between focal and recipient noses per trial, as a proxy for social interest of the pair, was measured during the interaction time, defined as the time that the two rats were simultaneously present in the choice area. Pairs with dominant focals showed lower nose-to-nose distance on both the first and second day of the PCT (independent sample t test for day 1: t_(20)_ = -2.11, p = 0.048, BF10 = 1.73; for day 2: t_(20)_ = -2.54, p = 0.02, BF10 = 3.18), suggesting that more proximal interactions preceded the emergence of prosocial choice (observed from day 2). Dynamics of nose-to-nose distance in the first seconds of interaction on (**B**) day 1 and (**C**) day 2. Pairs with dominant focal maintained shorter nose-to-nose distance in selfish trials on both days (one-way ANOVA comparing the four conditions for day 1: F_(3, 907)_ = 38.72, p = 1.5e-23; for day 2: F_(3, 1075)_ = 92.91, p = 1.825e-53). (**D**) The median distance between recipients’ nose and the central wall per trial was similar between submissive and dominant recipients on both the first and second day of the PCT when not taking into account trial type (independent sample t test for day 1: t_(20)_ = -1.21, p = 0.242, BF10 = 0.65; for day 2: t_(20)_ = -0.79, p = 0.442, BF10 =0.48). Nevertheless, the dynamics of the distance in the early phase of interaction on (**E**) day 1 and (**F**) day 2 showed that submissive recipients stayed closer to the wall than dominant recipients on day 2 in selfish trials, being this difference only marginal on day 1 (one-way ANOVA comparing the four conditions for day 1: F_(3,907)_ = 4.61, p = 0.003; for day 2: F_(3, 1075)_ = 4.84, p = 0.002). (**G**) The median nose speed per trial was similar between submissive and dominant recipients on both days of testing (independent samples t-test for day 1: t(20) = 0.47, p = 0.64, BF10 =0.42; for day 2: t(20)=0.05, p=0.96, BF10=0.39. Nevertheless, the dynamics of nose speed on (**H**) day 1 and (**I**) day 2 showed that submissive recipients kept moving the snout faster than dominant recipients on day 2 in selfish trials, and tended to do so on day 1, after the first second of interaction (one-way ANOVA comparing the four conditions for day 1: F_(3,884)_ = 3.29, p = 0.02; for day 2 F_(3,1037)_ = 6.63, p = 0.0002). (**J**) On both days, submissive recipients were more oriented towards their focal compared to dominant recipients (independent samples t test for day 1: t_(20)_ = -2.96, p = 0.008, BF10 =6.21; for day 2: t_(20)_ = -4.01, p = 0.0007, BF10=40.76). The same effect was observed in the dynamics of orientation (**K-L**), with submissive recipients being more oriented to their dominant decision-maker in selfish trials of both days. Interestingly on day 1, the orientation of submissive recipients in selfish trials was more similar to the orientation of dominant recipients in prosocial trials than that displayed in selfish trials. One-way ANOVA comparing the four conditions for day 1: F_(3,906)_ = 26.58, p = 1.76E-16; for day 2: F_(3,1075)_ = 76.92, p = 4.45E-45). **(M)** On both days, submissive recipients spent a higher proportion of time orienting towards their focal, while dominant recipients spent a higher proportion of time orienting away from their focal (for day 1, independent samples t test for the proportion of time with orientation <60°: t_(20)_ = 2.69, p = 0.015, BF10=3.8, time with orientation between 60° and 120°: t_(20)_ = -0.08, p = 0.939, BF10=0.39, time with orientation >120°: t_(20)_ = -3.08, p = 0.006, BF10=7.64 ; for day 2, time with orientation <60°: t_(20)_ = 2.49, p = 0.029, BF10=4.39, time with orientation between 60° and 120°: t_(20)_ = 2.26, p = 0.035, BF10=2.11, time with orientation >120°: t_(20)_ = -4.39, p = 0.0003, BF10 =82.7). Regarding mutual orientation, this trend was already significant on day 1 (on day 1, independent sample t test for the proportion of time with both rats’ orientations <60°: t_(20)_ = 2.249, p = 0.036, BF_10_ = 2.1, with both orientations between 60° and 120°: t_(20)_ = - 1.335, p = 0.197, BF_10_ = 0.73, with both orientations >120°: t_(20)_ = -2.654, p = 0.015, BF_10_ = 3.8); on day 2, time with both orientations <60°: t_(20)_ = 1.8, p = 0.087, BF_10_ = 1.12, with both orientations between 60° and 120°: t_(20)_ = 0.606, p = 0.551, BF_10_ = 0.44, with both orientations >120°: t_(20)_ = -3.274, p = 0.004, BF_10_ = 10.61). **(N)** Dominant focals investigate the wall that separates them from their submissive partner for longer durations regardless of trial type. Although this trend was not significant on day 1 due to the high variability observed (repeated measures ANOVA, “choice” (F_(1,20)_ = 2.29, p = 0.146, BFincl = 0.96), “choice” by “hierarchy” (F_(1,20)_ = 0.39, p = 0.541, BFincl =0.49) and “hierarchy” (F_(1,20)_ = 1.93, p=0.180, BFincl=0.74), it reached significant levels on day 2 ((repeated measures ANOVA, “choice” F_(1,20)_ = 0.12, p = 0.729, BFincl =0.3, “choice” by “hierarchy” F_(1,20)_ = 0.18, p = 0.672, BFincl=0.42, “hierarchy” F_(1,20)_ = 4.65, p = 0.043, BFincl =1.5). **(O)** Submissive and dominant recipients spent a similar amount of time investigating the wall when the focal was in the choice area, both in prosocial and selfish trials (RM-ANOVA for d1-2: “choice” (F_(1,20)_ = 2.14, p = 0.159, BFincl =0.69), “choice” by“hierarchy” (F_(1,20_) = 0.05, p = 0.827, BFincl=0.38) and “hierarchy” (F_(1,20)_ = 0.17, p=0.687, BFincl=0.47)). No effects were observed on the single days either (on day 1: “choice” F_(1,20)_ = 1.82, p = 0.193, BFincl =0.79, “choice” by “hierarchy” F_(1,20)_ = 1.28, p = 0.272, BFincl=0.63, “hierarchy” F_(1,20)_ = 0.0002, p = 0.989, BFincl=0.42; on day 2: “choice” F_(1,18)_ = 0.12, p = 0.739, BFincl =0.36, “choice” by “hierarchy” F_(1,18)_ = 2.35, p = 0.143, BFincl=1.05, “hierarchy” F_(1,18)_ = 0.93, p = 0.347, BFincl = 0.62. On **B,C,E,F,H,I,K,L** the left panel shows the temporal dynamics for each condition; the right panel shows the average, SEM and statistics of this time window. Letters show results of Student-Newman-Keuls (SNK) post-hoc test used to evaluate differences among dominance-trial type categories. *p<0.05, **p<0.01, ***p<0.001.

**Figure S4.**
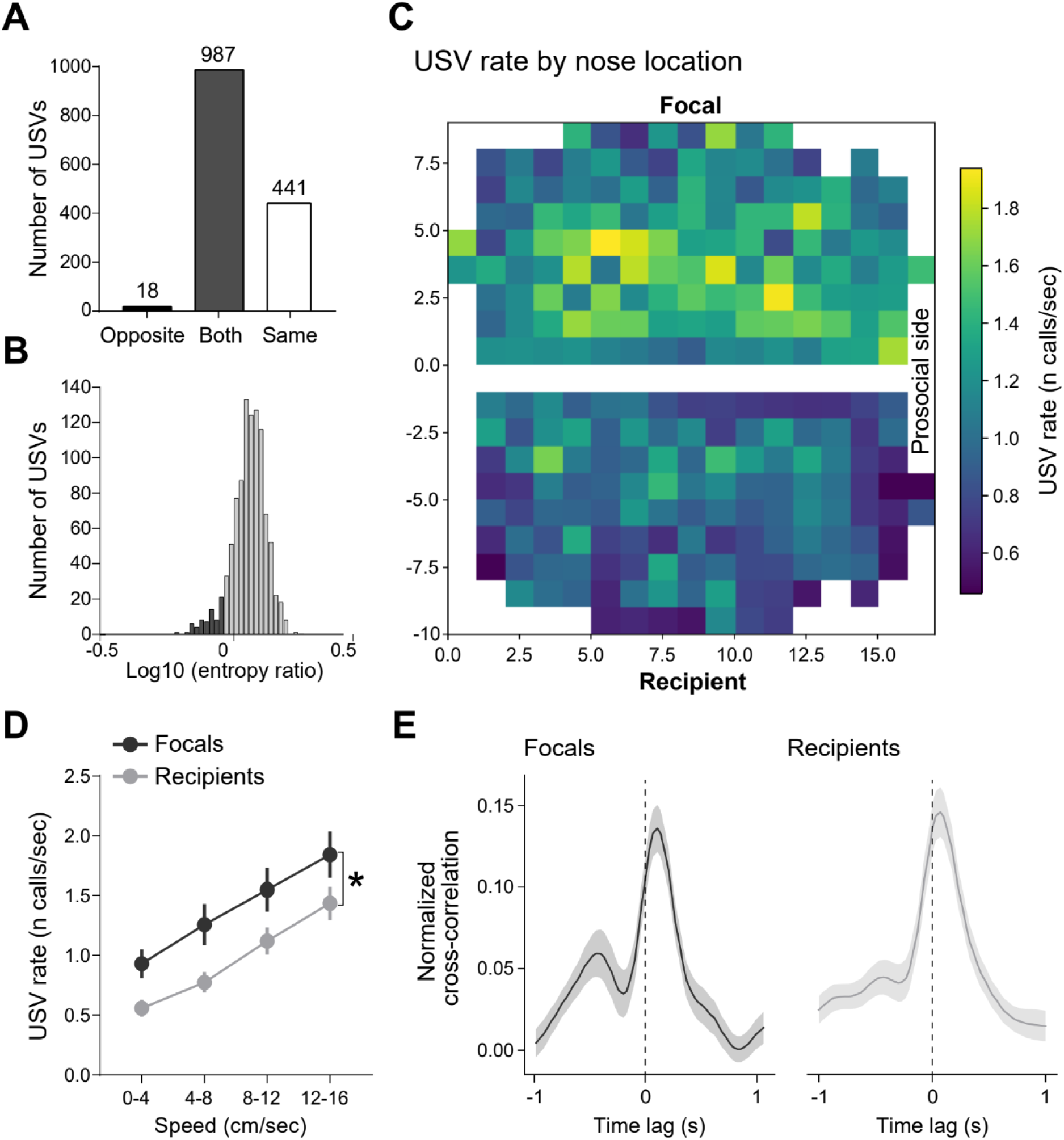
Validation of the USV assignment to individual animals within a pair, USVs rates by rat position and correlation between USV emission and snout speed. Related to Figure 5. **(A)** In recordings with only one rat placed alone in the double T-maze, all USVs should ideally be assigned to the side of the arena where the rat is. Of the 1446 USVs detected from 15 recordings obtained from 10 rats performing alone in one side of the double maze, 441 (31%) were only detected by the microphone over its side, 18 (1%) only by the opposite microphone and 987 (68%) were detected by both microphones. **(B)** Of those USVs detected by both microphones, 917 (93%) would have been correctly assigned to the one on the occupied side, as its signal had lower entropy (log_10_(entropy ratio occupied/empty) > 0, in grey). Overall, 94% of the detected calls were correctly assigned to the emitting rat (6% assignment error). **(C)** Heat map showing USVs rate of focal (top) and recipient rat (bottom) normalised by the time spent in each nose location of the choice area. Locations visited in less than 250 video frames were left out when normalising (white space). Prosocial side is on the right side of the image and dimensions are shown in centimetres. **(D)** USV rate vs instantaneous speed, showing that both focal and recipients increased call rate with speed. A RM-ANOVA indicated a significant effect of “speed bin” (F(1.7,71.68) = 130.26, p = 6e-23, ƞ^2^ = 0.195, BF_incl_ = 2e+35), no significant interaction between “speed bin” and “role” (F(1.7,71.68) = 0.47, p = 0.597, ƞ^2^ = 7e-4, BF_incl_ = 0.107) and a significant effect of “role” (F(1,42) = 4.87, p = 0.033, ƞ^2^ = 0.077, BF_incl_ = 1.48). *p<0.05. **(E)** Normalised cross-correlation of instantaneous speed and call rate for focals (left panel) and recipients (right panel). The peak of cross -correlation was consistently shifted from zero-time, revealing that vocal production preceded the speed increase by about 70ms. Mean across rats ± SEM are shown.

**Figure S5.**
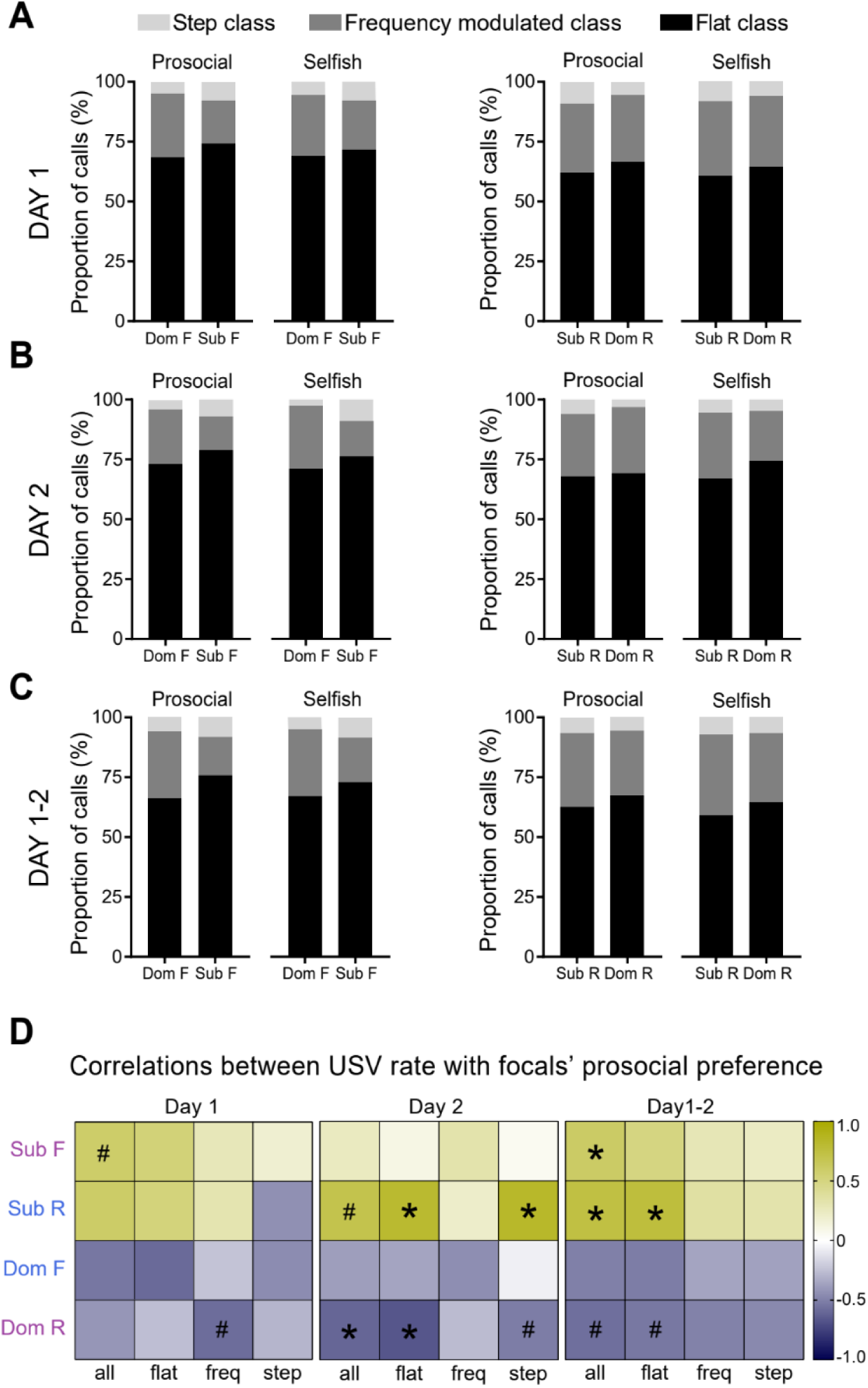
USV class proportions across days and correlation between USV rate and focal’s prosociality. Related to Figure 5. Proportion of USVs by class (flat, frequency modulated and step) on (**A**) day 1, (**B**) day 2 and (**C**) day1-2 in prosocial and selfish trials for both focal (left) and recipient (right) rats, according to hierarchy. Flat calls were the most frequent, followed by frequency modulated and step. No effect of dominance was observed in any condition, regardless of the day of testing and the role of the animals (repeated measure ANOVA of proportion of USVs class and trial type as within subjects factors. For focal rats: Class by hierarchy on day 1 F_(2,36)_=0.70 p = 0.501; day 2 F_(2,40)_=1.63 p = 0.208; day1-2 F_(2,40)_=1.157, p=0.325; Class by trial type and by hierarchy on day 1 F_(2,36)_=0.70 p = 0.502; day 2 F_(2,40)_=1.03 p = 0.365; day1-2 F_(2,40)_=0.163, p=0.85. For recipient rats: Class by hierarchy on day 1 F_(2,36)_=0.25 p = 0.78; day 2 F_(2,40)_=0.293 p = 0.748; day1-2 F_(2,40)_=0.355, p=0.703; Class by trial type and by hierarchy on day 1 F_(2,36)_=0.025 p = 0.976; day 2 F_(2,40)_=1.23 p = 0.304; day1-2 F_(2,40)_=1.561, p=0.222) (**D**) Partial correlations between USV rate and focals’ prosocial choice on day 1 (left panel), day 2 (middle panel) and day1-2 (right panel). Prosocial preference was positively correlated with recipient’s call rate only when the recipient was submissive (on day2 and day 1-2). Interestingly, a positive correlation was found also between prosociality of submissive focals and their own USV rate, only marginal on day 1 and significant on day 1-2, suggesting that increased prosociality is associated with increased call rate by submissive animals, regardless of their role in the task. In contrast, the USV rate of dominant recipients correlated negatively with prosociality by their submissive focals. This negative correlation was significant on day 2 and only marginal on day 1 and day 1-2. Dom F = Dominant Focal, Sub F = Submissive Focal, Sub R = Submissive Recipient, Dom R = Dominant Recipient. # p<0.1, * p<0.05.

**Figure S6.**
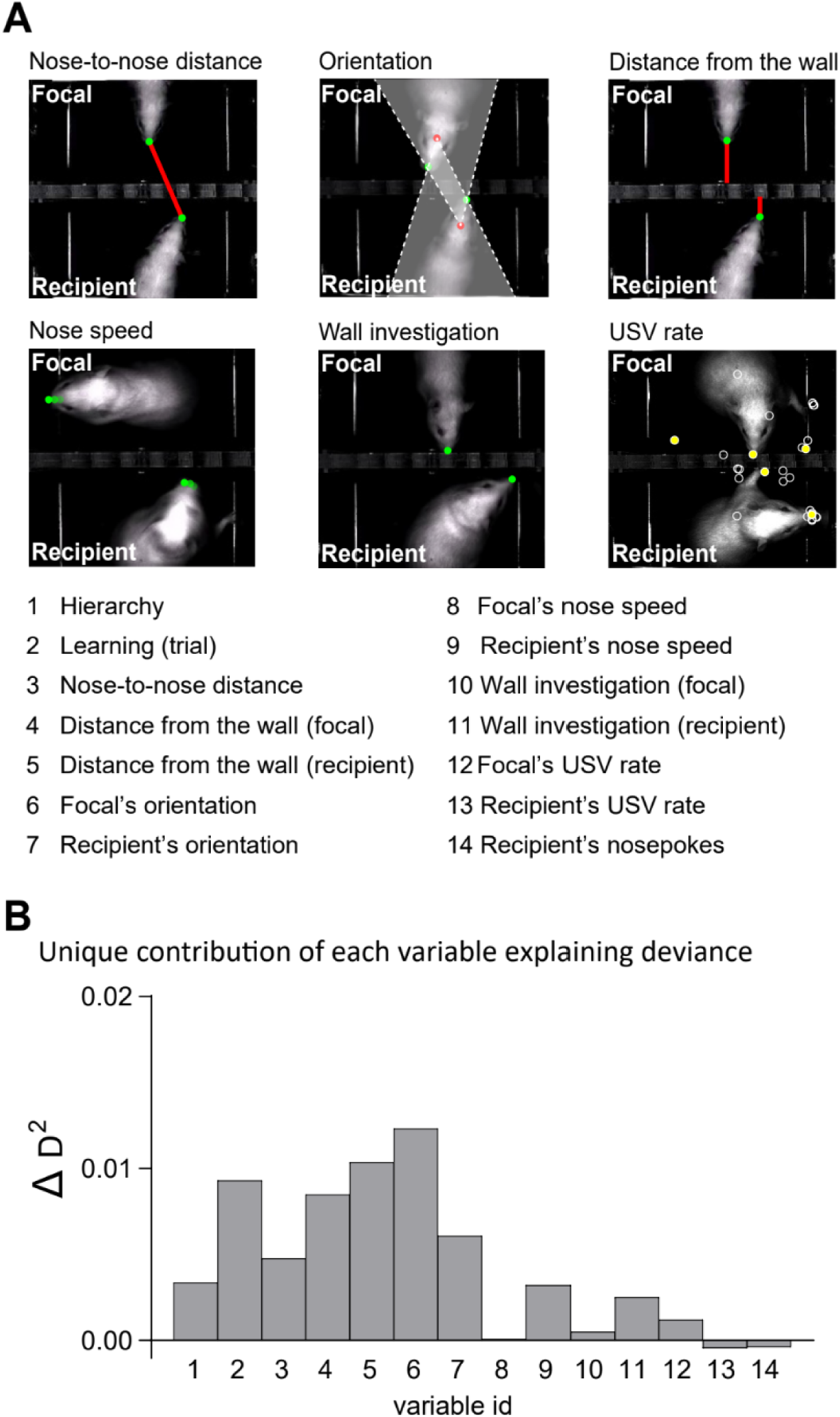
Unique contribution of each variable to the variability of the dataset. Related to Figure 6. (**A**) 14 behavioural and categorical parameters were chosen as regressors and evaluated by their contribution to the explained deviance of the model. Pictures represent the 14 behavioural variables measured either in focal and recipient animals, listed below. (**B**) The proportion of unique contribution (delta D^2^) of each regressor was very low, indicating that all variables were partially dependent on each other.

**Table S1.** Chance interval bounds generated by permutation test for each pair. Related to Figure S1. Low and high bounds show the 95% confidence interval for each focal animal. Dominant focals #: 3,7,8,9,10,12,15,19,46; Submissive focals #: 1,2,4,5,6,11,13,16,17,18,20,42,43; Familiar males #: 1,2,3,4,5,8,10,16,19,20,42,43,46; Unfamiliar males #: 21,22,23,24,25,26,27,28,29,30,31,41,44,45; Familiar females #: 47,48,49,50,51,52.

| Pair # | 1 | 2 | 3 | 4 | 5 | 6 | 7 | 8 | 9 | 10 | 11 | 12 | 13 | 14 | 15 |
| --- | --- | --- | --- | --- | --- | --- | --- | --- | --- | --- | --- | --- | --- | --- | --- |
| Low bound | -0,09 | -0,09 | -0,07 | -0,09 | -0,09 | -0,07 | -0,08 | -0,10 | -0,07 | -0,06 | -0,07 | -0,08 | -0,09 | -0,06 | -0,08 |
| High bound | 0,10 | 0,09 | 0,07 | 0,09 | 0,09 | 0,07 | 0,08 | 0,10 | 0,07 | 0,07 | 0,07 | 0,08 | 0,09 | 0,06 | 0,08 |
| Pair # | 16 | 17 | 18 | 19 | 20 | 21 | 22 | 23 | 24 | 25 | 26 | 27 | 28 | 29 | 30 |
| Low bound | -0,06 | -0,10 | -0,04 | -0,06 | -0,14 | -0,07 | -0,11 | -0,09 | -0,08 | -0,07 | -0,06 | -0,06 | -0,09 | -0,13 | -0,08 |
| High bound | 0,06 | 0,10 | 0,04 | 0,06 | 0,14 | 0,07 | 0,11 | 0,10 | 0,08 | 0,07 | 0,06 | 0,06 | 0,09 | 0,13 | 0,08 |
| Pair # | 31 | 41 | 42 | 43 | 44 | 45 | 46 | 47 | 48 | 49 | 50 | 51 | 52 |  |  |
| Low bound | -0,07 | -0,10 | -0,09 | -0,13 | -0,07 | -0,07 | -0,10 | -0,07 | -0,07 | -0,06 | -0,07 | -0,06 | -0,07 |  |  |
| High bound | 0,07 | 0,10 | 0,09 | 0,13 | 0,07 | 0,07 | 0,10 | 0,07 | 0,07 | 0,06 | 0,07 | 0,06 | 0,07 |  |  |

**Table S2.** Generalised Linear Model with all behavioural variables. Related to Figure 6. Prosocial choice is a binary variable (1:prosocial choice, 0:selfish choice). Trial is an ordinal variable indicating cumulative trial number over the first two days of the PCT. The dataset includes 1995 observations from 22 pairs of animals. *p<0.05, **p<0.01, *** p<0.001.

| Prosocial choice ~ trial * hierarchy * (nose-to-nose distance + focal's orientation + recipient's orientation) |  |  |  |  |
| --- | --- | --- | --- | --- |
| Variable | Estimate | z | p |  |
| (Intercept) | 2.771 | 6.234 | <0.00001 | *** |
| Trial | 0.033 | 3.748 | 0.00020 | *** |
| Hierarchy | -1.331 | -1.988 | 0.04690 | * |
| Nose-to-nose distance | 0.147 | 1.326 | 0.18480 |  |
| Focal's orientation | -0.033 | -4.469 | <0.00001 | *** |
| Recipient's orientation | -0.020 | -2.828 | 0.00470 | ** |
| Trial × Hierarchy | 0.049 | 3.268 | 0.00110 | ** |
| Trial × nose-to-nose distance | -0.004 | -1.572 | 0.11590 |  |
| Trial × focal's orientation | <0.001 | 1.013 | 0.31090 |  |
| Trial × recipient's orientation | <-0.001 | -0.193 | 0.84670 |  |
| Hierarchy × nose-to-nose distance | -0.449 | -2.639 | 0.00830 | ** |
| Hierarchy × focal's orientation | 0.032 | 2.945 | 0.00320 | ** |
| Hierarchy × recipient's orientation | 0.035 | 3.058 | 0.00220 | ** |
| Trial × hierarchy × nose-to-nose distance | 0.009 | 2.424 | 0.01530 | * |
| Trial × hierarchy × focal's orientation | -0.001 | -3.250 | 0.00120 | ** |
| Trial × hierarchy × recipient's orientation | -0.001 | -3.008 | 0.00260 | ** |

**Table S3.** Supplementary Table 3 | Reduced GLM: pairs with dominant focal. Related to Figure 6. Prosocial choice is a binary variable (1:prosocial choice, 0:selfish choice). Trial is an ordinal variable indicating cumulative trial number over the first two days of the PCT. The dataset includes 885 observations from 9 pairs of animals. #p<0.1, *p<0.05, **p<0.01, *** p<0.001.

| Prosocial choice ~ trial * (nose-to-nose distance + focal's orientation + recipient's orientation) |  |  |  |  |
| --- | --- | --- | --- | --- |
| Variable | Estimate | z | p |  |
| (Intercept) | 1.441 | 2.878 | 0.00400 | ** |
| Trial | 0.082 | 6.788 | <0.00001 | *** |
| Nose-to-nose distance | -0.302 | -2.340 | 0.01930 | * |
| Focal's orientation | -0.001 | -0.123 | 0.90210 |  |
| Recipient's orientation | 0.014 | 1.633 | 0.10250 |  |
| Trial × nose-to-nose distance | 0.005 | 1.848 | 0.06460 | # |
| Trial × focal's orientation | -0.001 | -3.358 | 0.00080 | *** |
| Trial × recipient's orientation | -0.001 | -3.817 | 0.00010 | *** |

**Table S4.** Reduced GLM: pairs with submissive focal. Related to Figure 6. Prosocial choice is a binary variable (1:prosocial choice, 0:selfish choice). Trial is an ordinal variable indicating cumulative trial number over the first two days of the PCT. The dataset includes 1110 observations from 13 pairs of animals. **p<0.01, *** p<0.001.

| Prosocial choice ~ trial * (nose-to-nose distance + focal's orientation + recipient's orientation) |  |  |  |  |
| --- | --- | --- | --- | --- |
| Variable | Estimate | z | p |  |
| (Intercept) | 2.771 | 6.234 | <0.00001 | *** |
| Trial | 0.033 | 3.749 | 0.00020 | *** |
| Nose-to-nose distance | 0.147 | 1.326 | 0.18480 |  |
| Focal's orientation | -0.033 | -4.469 | <0.00001 | *** |
| Recipient's orientation | -0.020 | -2.828 | 0.00470 | ** |
| Trial × nose-to-nose distance | -0.004 | -1.572 | 0.11590 |  |
| Trial × focal's orientation | <0.001 | 1.013 | 0.31090 |  |
| Trial × recipient's orientation | <-0.001 | -0.193 | 0.84660 |  |

